# Herbivore-induced activation of viral phosphatase disarms plant antiviral immunities for pathogen transmission

**DOI:** 10.1101/2020.06.17.158212

**Authors:** Pingzhi Zhao, Ning Wang, Xiangmei Yao, Changxiang Zhu, Saskia A. Hogenhout, Shu-sheng Liu, Xueping Zhou, Rongxiang Fang, Jian Ye

**Author notes:** These authors contributed equally to this work (Pingzhi Zhao, Ning Wang).

## Abstract

The survival of pathogens depends on their ability to overcome host immunity, especially arthropod-borne viruses (arboviruses) which must withstand the immune responses of both the host and the arthropod vector. Successful arboviruses often modify host immunity to accelerate pathogen transmission; however, few studies have explored the underlying mechanism. Here we report attracted herbivore infestation on the virus-infected plants promote transmission by the associated vector herbivore. This herbivore-induced defense suppression underpins a subversive mechanism used by *Begomovirus*, the largest genus of plant viruses, to compromise host defense for pathogen transmission. Begomovirus-infected plants accumulated βC1 proteins in the phloem where they were bound to host defense regulators, transcription factor WRKY20 and two mitogen-activated protein kinases MPK3 and MPK6. Once perceiving whitefly herbivory or endogenous secreted peptide PEP1, the plants started dephosphorylation on serine^33^ and stimulated βC1 protein as a phosphatase. βC1 dephosphorylated MPK3/6 and WRKY20, the latter negatively regulated salicylic acid signaling and vascular callose deposition. This viral hijacking of WRKY20 accumulated more vascular callose by which enforced whitefly prolonged salivation and phloem sap ingestion, therefore impelling more virus transmission among plants. We present a scenario in which viruses dynamically respond to the presence of their vectors, suppressing host immunity and promoting pathogen transmission only when needed.

## Introduction

In nature ecosystems, majority of emerged or reemerged pandemic diseases are caused by zoonotic or arthropod-borne viruses (arboviruses), e.g. Zika virus and begomoviruses (Weaver et al., 2018; Gnanasekaran et al., 2019). Being obligate parasites, viruses usually depend for survival on being able to spread. Only if they can spread from host to host rapidly in community, viruses are then important economically. The transmission potential of pathogens is one of the most important virulence factors that cause the outbreak and epidemic of diseases, such as COVID-19 caused by SARS-associated novel coronavirus 2, SARS-CoV-2 (Li et al., 2020). Arboviruses commonly modify the behavior of the arthropod vector in a manner that facilitates pathogen transmission (Eigenbrode et al., 2018; Weaver et al., 2018). For instance, mosquito-borne bunyaviruses and flaviviruses as well as thrip-borne tospoviruses promote virus transmission by directly increasing vector biting rates and by interfering with the ingestion process (Stafford et al., 2011; Liu et al., 2017). Furthermore, arboviruses may indirectly promote transmission by 1) manipulating the immune responses of both the vector and the host, 2) suppressing host immunity by vector salivary factors, 3) manipulating vector behavior, and 4) encoding transmission facilitators (Vasilakis and Tesh 2015; Wang et al., 2019; Wu et al. 2019; Wu and Ye, 2020). Since plants are sessile, around 80% plant viruses are transmitted by insect vectors, especially whiteflies and aphids that transmit more than 50% of all vector-borne viruses. Their transmission depends on multilayered virus–vector–host interactions that are associated with changes in vector feeding behaviors and with alterations in plant host’s morphology and/or metabolism to favor the attraction or deterrence of vectors (Eigenbrode et al., 2018; Mauck et al., 2019).

Plant viruses are masters at reprogramming defensive traits and other host processes. The genus *Begomovirus* (family *Geminiviridae*) is believed to cause the earliest known plant virus disease over a millennium ago and now consists of more than 440 species and represents the largest and most diverse genus of plant viruses (Yang et al., 2019). Exclusively transmitted by insect vector whitefly (*Bemisia tabaci*), begomoviruses cause several devastating crop diseases (Liu et al., 2007; Yang et al., 2019). Cotton leaf curl virus and tomato leaf curl virus, are two clusters of invasive monopartite begomoviruses, which are always associated with β-satellite (Zhao et al., 2019). We and other groups have demonstrated that the βC1 protein encoded by their associated β-satellites suppresses host antiviral and anti-herbivore defense (Jia et al., 2016; Yang et al., 2019; Zhang et al., 2012; Li et al., 2014). We have recently shown that begomoviral βC1 mobilizes host immunity to deter nonvector herbivores, suggesting complex immune network among this tripartite interaction of virus-plant-herbivore (Zhao et al., 2019). The interactions between host immunopathological responses and co-infections from begomovirus plus herbivores, however, have remained obscure, as well as its potential biological relevance.

Host plants have evolved many defensive mechanisms to circumvent against the invasion of pathogens and pests including gene silencing, phytohormones and innate immunity. A few of molecular patterns and their corresponding receptors have been identified one by one and shown to elicit a set of downstream molecular responses (Saijo et al., 2018; Erb and Reymond, 2019). By perceiving exogenous stresses, e.g. herbivore or pathogen attacks, plant activates a variety of early defense responses including kinase activation such as mitogen-activated protein kinase (MAPK) and sucrose non-fermenting 1 related protein kinase (SnRK), calcium (Ca^2+^) influxes, and defense hormones signaling (Schuman and Baldwin, 2016; Zhou et al., 2019). Plants reprogram their downstream transcriptional network composing with AP2/ERF, MYB, bHLH and WRKY families to regulate biosynthesis of specialized metabolites such as terpenoids, phytoalexins, reactive oxygen species (ROS) burst including hydrogen peroxide (H_2_O_2_) and callose deposition (Tsuda and Somssich, 2015). Phytohormone salicylic acid (SA) induces deposition of callose at the cellular sites of attack (Li et al., 2017; Huang et al., 2019). Callose is a β(1,3) glucan polymer that provides focused delivery of chemical defenses. However, whether and how the begomovirus or/and whitefly counter these host defenses to provide pro-viral infectivity and transmission are unclear.

In this study, we report a phenomenon of herbivore-induced defense suppression which promotes virus transmission among plants. Once perceiving whitefly herbivory by PEPR1/2 receptors, begomovirus-infected plants activate dephosphorylation on serine^33^ of the begomoviral βC1 through suppression of plant SnRKs and stimulate βC1 to act as a phosphatase. βC1 directly deactivates mitogen-activated protein kinases MPK3/6 and the vascular transcription factor WRKY20. This begomoviral hijacking of plant phosphorylation cascade activates host salicylic acid signaling and enhances plant vascular callose deposition, which imposes changes of whitefly feeding behavior, leading to more virus transmission among plants. Our results have broad implications for antiviral through genetic or chemical interference with vector transmission.

## Results

### Herbivore-induced defense suppression in begomovirus-infected plants to promote virus transmission

To compare host defense responses between single stress and multiple stresses, we investigated the defense responses upon herbivory attack on the cotton leaf curl Multan virus (CLCuMuV) complex (CA+Cβ)-infected *Gossypium barbadense* (cotton) using uninfected cotton plants as control. Phosphorylation levels of MAPKs indicate active plant immunity. Figure 1A shows that plant immunity could rapidly activate response to single infection from vascular feeding whitefly and the green peach aphid (*Myzus persicae*), leaf-chewing herbivore cotton bollworm (CBM; *Helicoverpa armigera*) and the viral pathogen. Aphid is another piercing insect vector and transmits over 300 plant viruses (Hooks and Fereres, 2006). As expected in most cases of double combined stresses, the immune responses have addictive effect and MAPKs phosphorylation is considerable higher than single stress. Unexpectedly, the presence of whitefly did not activate but reduced the active immunity that induced by the CLCuMuV or cotton bollworm.

**Figure 1.**
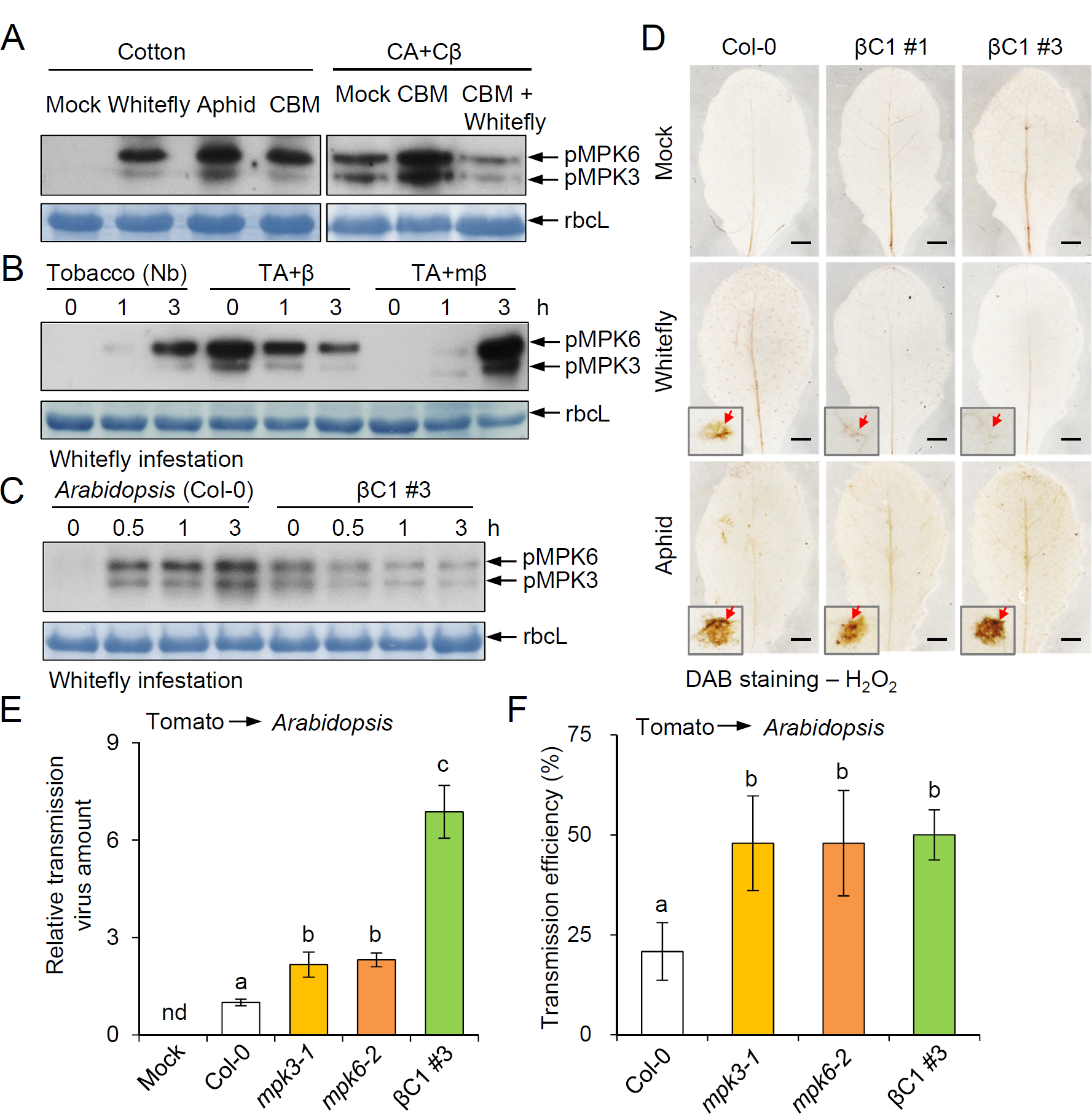
Whitefly-induced defense suppression promotes begomovirus transmission between plants. (**A**) MAP kinase activation in uninfected cotton and the CLCuMuV complex (CA+Cβ)–infected cotton upon herbivore infestation. (**B**) MAP kinase activation in uninfected tobacco (Nb) and the TYLCCNV complex (TA+β)–infected or βC1 betasatellite mutant (TA+mβ) Nb plants upon whitefly infestation for indicated time. (**C**) MAP kinase activation in transgenic *Arabidopsis* plants ectopically expressing *βC1* upon whitefly infestation for the indicated periods. In **A**-**C**, phosphorylation levels of MAPKs were detected by immunoblotting using anti-phospho-p44/42 antibody. Stained membrane bands of the large subunit of Rubisco (rbcL) were used as a loading control. (**D**) H_2_O_2_ level with or without herbivore treatment on wild-type Col-0 plants and *βC1-*expression *Arabidopsis* lines. Red arrows in insets indicate the amplified feeding sites of whitefly or aphid on plant leaves. Scale bars = 2 mm. (**E**) The relative TYLCV virus amount transmitted from tomato to *Arabidopsis* by whiteflies. Col-0 plants infestated with non-viruliferous whiteflies were used as the mock. Bars represent means ± SD (n=8). (**F**) Virus transmission efficiency from tomato to Col-0 or *mpk3-1, mpk6*-2 mutant or *βC1-*expression *Arabidopsis* by whiteflies. The efficiency represents as the percentage of virus-positive plants against total 8 challenged plants as a repeat. Three biological repeats were taken and results were represented as means ± SD (n= 3). In **E**,**F**, lowercase letters indicate significant differences among different lines according to one-way ANOVA followed by Duncan’s multiple range test (P<0.05).

As most monopartite begomoviruses are associated with β-satellite, we next tested whether this herbivore-induced defense suppression (HIDS) also occurs in model plant hosts which could be infected by another β-satellite–associated begomovirus. *Nicotiana benthamiana* (tobacco, Nb) plants infected with tomato yellow leaf curl China virus (TYLCCNV) and its associated β-satellite (TA+β), had a similar HIDS effect upon the infestation of vector whitefly (Figure 1B and Figure 1-figure supplement 1A,B). This HIDS also occurred in another model plant *Arabidopsis thaliana*, whose immunities and defense pathways are well understood. Most of defense genes associated with pattern-recognition receptor-mediated pathogen perception and signal transduction in TA+β-infected plants were suppressed by the infestation of whitefly (Figure 1-figure supplement 1C,D). Neither another RNA virus, <u>tobacco rattle virus</u> (TRV) nor a begomovirus nonvector green peach aphid, elicited the HIDS (Figure 1-figure supplement 2A,B). Dead whitefly also could not trigger this HIDS (Figure 1-figure supplement 2C). These results demonstrate that this HIDS is specifically induced in response to begomovirus infection and whitefly feeding.

Two opposing hypotheses may be proposed to explain the herbivory-induced host defense suppression. One is that the host protein may be repressed e.g. via salivary effector Bsp9 of whitefly that shuts down host defense system reported previously (Wang et al., 2019). Since whitefly herbivory itself can activate plant defense instead of suppression, this explanation seems most unlikely. The other is that the herbivore induction may underpin a subversive mechanism used by the virus to compromise host defense. Previously we demonstrated that βC1 protein encoded by begomovirus could suppress host defenses. We next tested whether this HIDS is dependent on the βC1 protein. As shown in Figure 1B-D and Figure 1-figure supplement 3, HIDS could reproduce in two transgenic lines expressing *βC1* (βC1 #1 and βC1 #3 lines) plants but failed in the TYLCCNV/TYLCCNB βC1 mutant (TA+mβ)-infected plants upon whitefly infestation, suggesting that βC1 is the viral determinator and necessary for whitefly-induced defense suppression.

To investigate the potential biological relevance of this HIDS, we checked the final consequence of this herbivore behavior on virus-infected plants. We quantified virus infectivity and quantity of virus transmitted by whiteflies from *Solanum lycopersicum* (tomato) to *Arabidopsis*. To avoid a possible direct interference of whitefly feeding behavior by βC1 protein, we used whiteflies carrying the closely related begomovirus tomato yellow leaf curl virus (TYLCV) in a limited transmission period (72 h) as we demonstrated before (Wang et al., 2019). The quantity of virus in the βC1-plants and two *mpk* mutants was about 10-fold and 2-fold higher than in wild-type Col-0 plants (Figure 1E). Furthermore, virus infectivity was also much higher on two *mpk* mutants and βC1-plants (about 50%) than on wild-type plants (20.8%) at 15 days post whitefly infestation (Figure 1F). These results strongly indicate that this begomoviral βC1– involved HIDS promotes virus transmission and infectivity among plants.

### βC1 phosphatase targets and inactivates plant MAPKs

To identify how βC1 controls this HIDS effect, we sought to identify βC1-targeted host factors. Because βC1-whitefly paired could suppress MAPKs activation and MAPK-related immune signaling often involves protein-protein interactions, we raised the possibility that the viral βC1 acts directly on components of the MAPK cascade. We found that the βC1 directly interacts with *Arabidopsis* MPK3 and MPK6 in yeast two-hybrid assay (Figure 2A). We confirmed the interactions between *Arabidopsis* MPK3/MPK6 and βC1 by *in vitro* GST-pull down and *in vivo* bimolecular fluorescence complementation (BiFC), and co-immunoprecipitation (CoIP) assays (Figure 2B,C and Figure 2-figure supplement 1). Considering the reduced phosphorylation levels of MPK3/6 in βC1 transgenic plants upon whitefly infestation (Figure 1C and Figure 1-figure supplement 3), we tested whether βC1 directly affects the phosphorylation levels of MPK3 and MPK6. We used recombinant His-MKK5^DD^ (threonine 215 and serine 221 mutated to aspartate, a constitutively active form) to phosphorylate purified His-MPK6 protein (Zhang et al., 2007). Inclusion of different amount of purified βC1 protein in the reactions substantially reduced the phosphorylation levels of MPK6 (Figure 2D), suggesting that βC1 inactivates plant MAPKs presumably by dephosphorylation.

**Figure 2.**
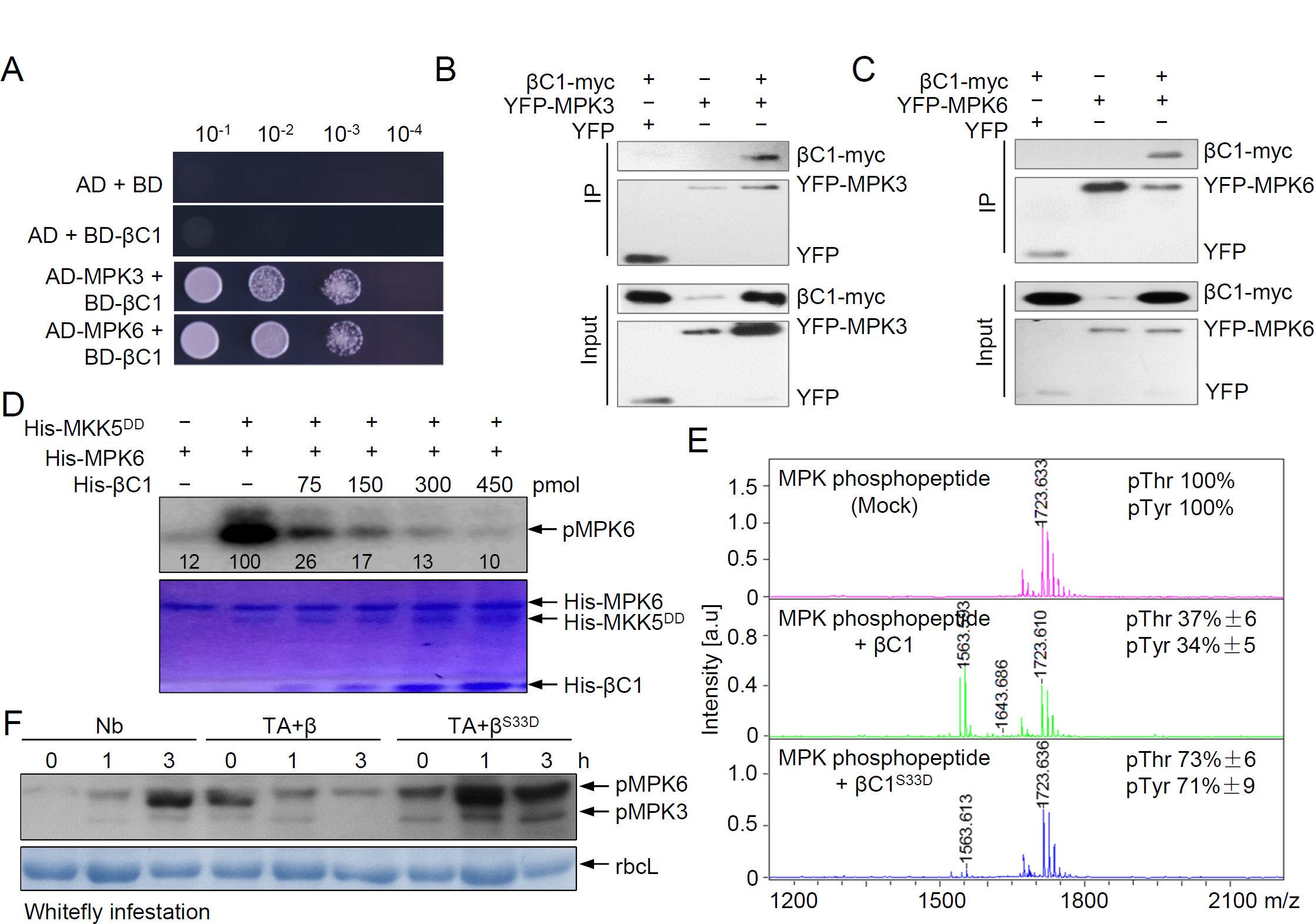
βC1 interacts with and dephosphorylates *Arabidopsis* MPK6. (**A**) βC1 interaction with MPK3 or MPK6 in a yeast two-hybrid assay. The empty vectors pGAD424 and pGBT9 were used as negative controls. (**B, C**) Co-IP analysis of the interaction between YFP-MPK3 (**B**) or YFP-MPK6 (**C**) and βC1-myc *in vivo*. Samples before (Input) and after immunoprecipitation (IP) were analyzed by immunoblot analysis using monoclonal antibody against YFP or myc. (**D**) βC1 dephosphorylates MPK6 *in vitro*. Phosphorylation levels of pMPK6 were estimated by band intensities in immunoblots and further normalized by loaded protein amount indicated by coomassie brilliant blue staining using ImageJ software. (**E**) *In vitro* phosphatase activity of βC1 and the phosphate mimic mutant βC1^S33D^. The intensity of the phospho-Thr form or phospho-Tyr form of treated synthetic phosphopeptide in the presence of βC1 and βC1^S33D^ were calculated as each peak area in tandem mass spectrometry. The untreated (mock) synthetic phosphopeptide was set to 100%. Data represent mean ± SD (n=3). (**F**) MAP kinase activation upon whitefly infestation in Nb plants and plants infected by TA+β, the phosphate βC1 mutant virus TA+β^S33D^, or TA+mβ.

Plant MAPK is activated by the dual phosphorylation of threonine (Thr) and tyrosine (Tyr) residues in a conserved TXY motif (Hazzalin and Mahadevan, 2002). Thus, to further investigate the enzymatic activity of βC1 in dephosphorylating MAPKs, we synthesized an MPK6 phosphopeptide (SESDFM-pT-E-pYVVTR) carrying p-threonine (pT) and p-tyrosine (pY) at the TXY motif. Treatment with βC1 reduced the mass of the MPK6 phosphopeptide by 80 Da or 160 Da, as determined by matrix-assisted laser desorption/ionization time-of-flight mass spectrometry (MALDI-TOF-MS) (Figure 2E). The mass reduction of one or two phosphate groups from the TXY motif was confirmed by tandem mass spectrometry (Figure 2-figure supplement 2). Thus, βC1 protein specifically cleaves the CO-P bond of the phosphopeptide and may function like a typical protein phosphatase to inactivate MAPK.

We previously demonstrated that βC1 was phosphorylated by tomato SnRK1 mainly on the serine residue at position 33, and the phosphate βC1 mutant virus TA+β^S33D^ (serine 33 to aspartate)-infected plants reduced virus accumulation and *βC1* expression (Figure 2-figure supplement 3A,B) (Shen et al., 2011; Zhong et al., 2017). Serine^33^ residue mutant in the βC1 protein did not affect the interactions with MPK3 and MPK6 in the subcellular localization (Figure 2-figure supplement 3C). However, the βC1^S33D^ impaired its phosphatase activity (Figure 2E). The βC1 phosphatase activity is required for HIDS, as indicated by the lack of reduction of MPK3/6 (another protein name as SIPK/WIPK) activation in TA+β^S33D^-infected *N. benthamiana* plants upon whitefly infestation (Figure 2F and Figure 2-figure supplement 3D,E). There might be no internal relationship between this phosphatase activity and its previously known function as a RNA silencing suppressor for βC1, likely due to that βC1^S33D^ mutant did not decrease the activity as a RNA silencing suppressor compared with βC1 protein (Supplementary Figure 1). Therefore, phosphatase activity of βC1 protein might be independent for its known silencing suppressor activity.

### PEPR receptors involve in perception of whitefly infestation

So far we have demonstrated that begomoviral βC1 protein could compromise host defense. But it is still unknown how whitefly infestation underpins this MAPK subversive mechanism by begomoviruses. Host surveillance system perceives exogenous invasion by multiple membrane-localized Pattern Recognition Receptors (PRR), which will then trigger the activation of MAPKs and downstream defense signaling (Choi and Klessig, 2016). We noticed that expression of several PRR genes, including PEPR family, were greatly repressed by whitefly infestation in begomovirus-infected *Arabidopsis* (Figure 1-figure supplement 1D). The leucine-rich repeat protein kinase PEP RECEPTORs (PEPR1/PEPR2) are believed to sense the damage caused by herbivore feeding on plant cell (Huffaker et al., 2006). Besides, we observed that the expression of *Arabidopsis PEPR1* was induced by whitefly feeding (Supplementary Figure 2A) or by the presence of *Arabidopsis* Pep1, a 23-amino acid endogenous peptide (Wang et al., 2019). The facts that *Arabidopsis* PEPR1/PEPR2 is highly expressed in plant vascular (Bartels et al., 2013), where whiteflies and begomoviruses infect (Yang et al., 2019), prompts us to hypothesize that PEPRs are one major perception machinery to sense whitefly infestation for plant cell. We next asked whether this *PEPR1* expression induction is specific for whitefly, another vascular-feeding herbivore aphid was further tested. No significant gene expression induction upon aphid infestation was observed in Col-0 plants (Supplementary Figure 2B), suggesting that PEPR signaling may be specifically involved in the perception of infestation of whitefly due to their unique feeding apparatus and unique feeding behaviors. Furthermore, whitefly-induced MAPK activation was remarkably reduced in PEPR1/PEPR2 receptors mutant plants (*pepr1*/*pepr2*) compared with Col-0 plants (Supplementary Figure 2C), further indicating that PEPR1/PEPR2 are two of the receptors for sensing whitefly herbivory. Similar to whitefly infestation, Pep1-induced MAPK activity suppression was also observed in *βC1*-expression lines (Supplementary Figure 2D), likewise indicating that the βC1-mediated HIDS could be triggered by Pep1-PEPR1/PEPR2 recognition event. As no transcriptional suppression on *PEPR1* in βC1-expression plants occurred (Supplementary Figure 2A), this HIDS could be perceived via PEPR1/2 and MAPK phosphorylation cascade signaling in *Arabidopsis*.

### Whitefly herbivory or Pep1-treatment may attenuate SnRK-mediated phosphorylation on βC1 protein

If βC1 is a real protein phosphatase, it is difficult to explain why MAPK phosphorylation cascade could be highly activated in plants affected by a single stress, e.g. begomovirus infection. A hypothesis put forward by us above that herbivore induction may release a repressive modification on βC1 protein, could potentially offer a resolution of this contradiction. SnRK kinase families are associated with this repressive modification on βC1 protein. SnRKs involve in plant responses to several biotic stresses, including geminiviruses (Shen et al., 2011; Wurzinger et al., 2018). We have shown that SnRK proteins confer plant resistance to geminiviruses by phosphorylation modification on βC1 and symptom determinant. That particular high reduction in most of the defense genes’ transcripts upon whitefly infestation in begomovirus-infected plants (Figure 1-figure supplement 1D) prompts us to further hypothesize that SnRK family genes may play roles in the HIDS. To test this hypothesis, we conducted gene expression analysis for these *SnRK* family defense genes in begomovirus-infection and uninfected plants upon whitefly infestation or Pep1-treatment. Consistent with our hypothesis, the expression of nine of the tested ten *SnRK* genes was significantly reduced in begomovirus-infected plants upon whitefly infestation compare with those of uninfestated mock plants (Figure 3A and Figure 3-figure supplement 1A). The Pep1-treatment resulted in even much stronger effect of reduction on *SnRK* genes expression than those of whitefly-infestated plants. If the expression of these *SnRK* genes was reduced, we would expected that the repression of phosphorylation of βC1 protein would be alleviated upon whitefly herbivory. As we expected, phosphorylation levels of βC1 protein was dramatically and gradually reduced by whitefly herbivore or Pep1-treatment in *N. benthamiana* plants (Figure 3B). This whitefly-triggered reduction of phosphorylation level of βC1 seems be specific to the vector insect, since nonvector aphid infestation did not have any effect on the phosphorylation level (Figure 3-figure supplement 1B).

**Figure 3.**
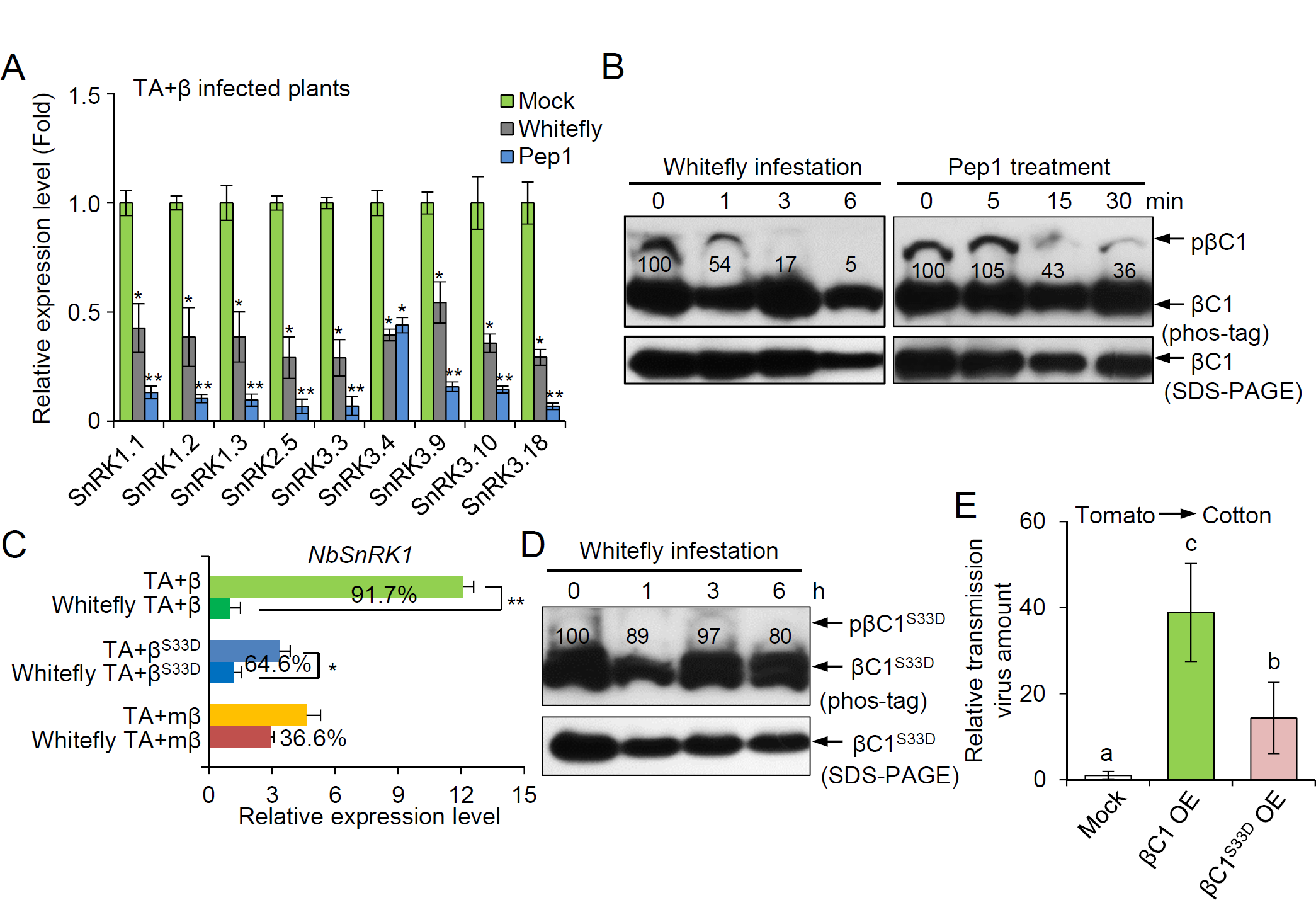
Herbivore-induced hypophosphorylation of βC1 in plants enhances virus transmission between plants. (**A**) Relative expression levels of *Arabidopsis SnRKs* upon whitefly infestation or Pep1 treatment in TA+β-infected Col-0 plants, which were compared with those in untreated TA+β-infected plants (Mock). Bars represent means ± SD (n= 3) (*, P< 0.05; **, P< 0.01; Student’s *t*-tests). (**B**) Levels of phosphorylated βC1 protein in transiently expressing *myc-βC1* Nb plants upon whitefly infestation or Pep1 treatment. Phosphorylated βC1 (pβC1) and total βC1 protein were quantified by band intensities in immunoblots using ImageJ software. The relative level of phosphorylation was normalized against the amount of each treatment at 0 h and shown below each band. (**C**) Relative expression levels of *NbSnRK1* in Nb plants infected by TA+β, TA+β^S33D^, or TA+mβ without or with whitefly infestation for 6 h. Bars represent means ± SD (n= 3) (*, P< 0.05; **, P< 0.01; Student’s *t*-tests). (**D**) Phosphorylation levels of the mutant βC1^S33D^ protein in transiently expressing *myc-βC1*^*S33D*^ Nb plants upon whitefly infestation for the indicated periods (hours). The relative level of phosphorylation was normalized against the amount of each treatment at 0 h. (**E**) The relative TYLCV virus amount transmitted from tomato to transiently expressing βC1 or βC1^S33D^ protein cotton by whiteflies. Cottons infestated with non-viruliferous whiteflies were used as the mock. Bars represent means ± SD (n=8). Lowercase letters indicate significant differences among different lines according to one-way ANOVA followed by Duncan’s multiple range test (P < 0.05).

This SnRK-involved HIDS could also be observed in another begomovirus-infected plant *N. benthamiana* but much dampened in phosphate-defected βC1^S33D^ mutant virus. We found that *NbSnRKs* expression in TA+β^S33D^-infected *N. benthamiana* plants were decreased upon whitefly infestation, but to a lesser extent than in TA+β-infected plants (Figure 3C and Figure 3-figure supplement 1C). Consistently, whitefly infestation had little impact on the phosphorylation level of βC1^S33D^ protein (Figure 3D). Meanwhile, we speculated that more phosphorylation level of βC1 would highly dampen its roles in promoting virus transmission. We further found that the constitutive phosphorylation on serine^33^ of βC1 much decreased the quantity of virus transmitted; the quantity in the mutant was only 37% of that in the wild type βC1 protein (Figure 3E). Together, these results show that the serine^33^ of βC1 is essential for evading host SnRK-mediated defense and contributes to enhance virus transmission.

### HIDS promotes begomovirus accumulation in plants

The SnRK and MPKs are kinases to amplify host defense responses to counter pathogen invasion. The suppression of MAPK activation and *SnRK* gene expression suggested that βC1 targets an early step of plant defenses. We thus tested the effect of βC1 on two other known signaling events: Ca^2+^ flux, and ROS burst. These two defense responses can be induced by biotic stresses such as plant pathogens and herbivores. Here we used Pep1 treatment to mimic whitefly herbivory. To monitor Ca^2+^ levels within the cell over time, we crossed Col-0 *Arabidopsis* expressing GFP-based cytoplasmic Ca^2+^ sensor (*GCaMP3*) (Vincent et al., 2017) with two *mpk* mutants, *pepr1/2* mutant and *βC1* overexpression plants. The Pep1-triggered cytoplasmic Ca^2+^ elevation is PEPR-dependent, since this response was almost abolished in *pepr1/2*/*GCaMP3* mutant (Figure 4A). As expected, βC1 strongly suppressed the Pep1-triggered Ca^2+^ elevation, partially phenocopying *MAPK*-deficient mutants. ROS is a key host free radical-defense against biotic stresses (Guo et al., 2018). ROS accumulation was suppressed by whitefly infestation on βC1-plants, as indicated by tissue staining using 3.3′-diaminobenzidine (DAB) (Figure 1D). Similar to the effect of whitefly infestation, upon induction of Pep1, the βC1-plants showed a reduction in ROS accumulation, compared with the induced strong accumulation of ROS in tissues of the wild-type Col-0 plants (Figure 4B).

**Figure 4.**
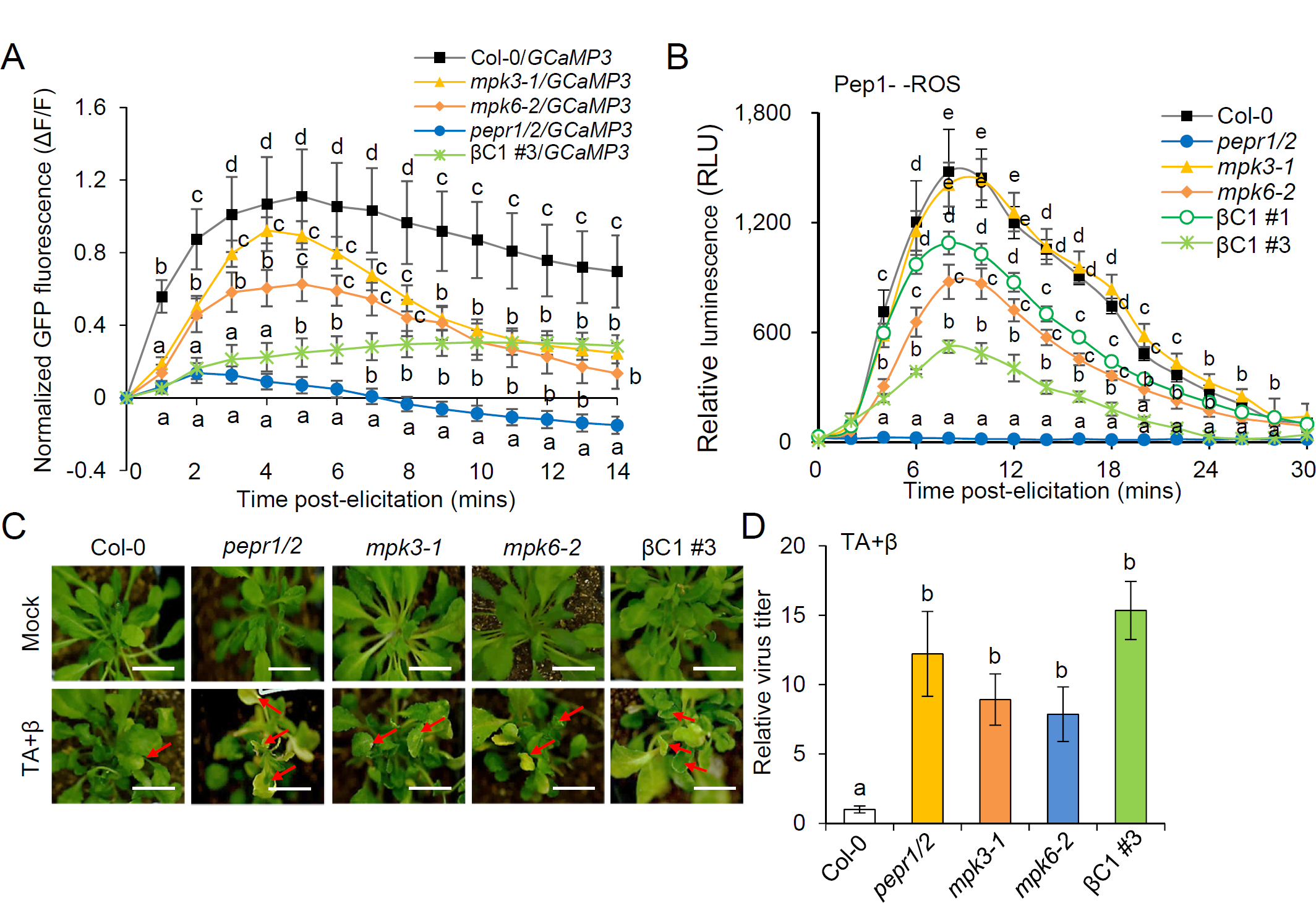
Pep1-induced defense suppression promotes begomovirus accumulation. (**A**) The Pep1-eliciting cytoplasmic Ca^2+^ elevations in *35S: GCaMP* plants crossing with either *βC1-*expression or *mpk3-1* or *mpk6-2* or *pepr1/2* plants. Normalized GFP fluorescence (ΔF/F) represents cytoplasmic Ca^2+^ elevations within 14 min of Pep1 treatment. F, average fluorescence intensity at the first record (baseline); ΔF, difference between measured fluorescence and baseline fluorescence. Bars represent means ± SD (n=15). (**B**) ROS production induced by Pep1 treatment was monitored for 30 min using a luminol-based assay in leaf discs derived from different genotypes of *Arabidopsis*. Data are given as relative luminescence units (RLU). Bars represent means ± SD (n=16). (**C**) Typical symptoms of TA+β-infected *Arabidopsis* plants at 21 dpi as indicated by red arrows. Scale bars = 1 cm. (**D**) Relative virus titer accumulated in TA+β-infected *Arabidopsis* plants at 21 dpi. Bars represent means ± SD (n=8). Lowercase letters indicate significant differences among different lines according to one-way ANOVA followed by Duncan’s multiple range test (P < 0.05).

Both Ca^2+^ homeostasis and ROS production are widely proven to play important roles in defense against invading pathogens and prevent their further proliferation in cells (Flury et al., 2013; Chen et al., 2019). If these known host defense responses have been shut down by βC1, then we would expect that host resistance against the begomovirus is compromised. To test this hypothesis, we inoculated these mutant plants with the TA+β virus in *Arabidopsis*. Twenty one days after inoculation with TA+β, *pepr1/2* mutant, two *mpk* mutant and βC1-plants showed more strongly curled leaves and higher virus accumulation compared to the wild type Col-0 plants (Figure 4C,D). These results together indicate that HIDS promotes viral pathogenesis.

### βC1 impairs the phosphorylation cascade of MAPK-WRKY20

So far, we have demonstrated the early defense signaling of this HIDS in begomovirus-infected host plants. It is still unclear how the virus transmission is affected under this circumstance. To this end, we continued to ask what’s the direct and downstream molecular or protein responsible for enhancing virus transmission among plants. Recently, we have showed that the βC1 interferes with plant WRKY20 to change plant-herbivore interactions (Zhao et al., 2019). Since the PEPR1/2-MAPK cascade specifically recognizes whitefly herbivory in begomovirus-infected plants, thus we speculated that WRKY20 is one of the MAPK direct substrates to reprogram plant defense and affect whitefly-mediated virus transmission. We thus tested the hypothesis by conducting yeast two-hybrid and BiFC assays, and confirmed that both MPK3 and MPK6 physically interact with WRKY20 (Figure 5-figure supplement 1). Since βC1 could interacts with any of MPK3/MPK6/WRKY20, we raised the possibility that βC1 competes with MPK3 and MPK6 for interaction with WRKY20. To test this hypothesis, we performed a modified BiFC assay. A negative control of GUS coexpression did not affect the interaction between MPK3 or MPK6 and WRKY20; whereas coexpressing with βC1, the interaction signal strength of MPK3-WRKY20 or MPK6-WRKY20 indicated by EYFP fluorescence intensity markedly decreased (Figure 5-figure supplement 2). The results indicate that βC1 interferes with the interaction between MPK3 or MPK6 and WRKY20.

**Figure 5.**
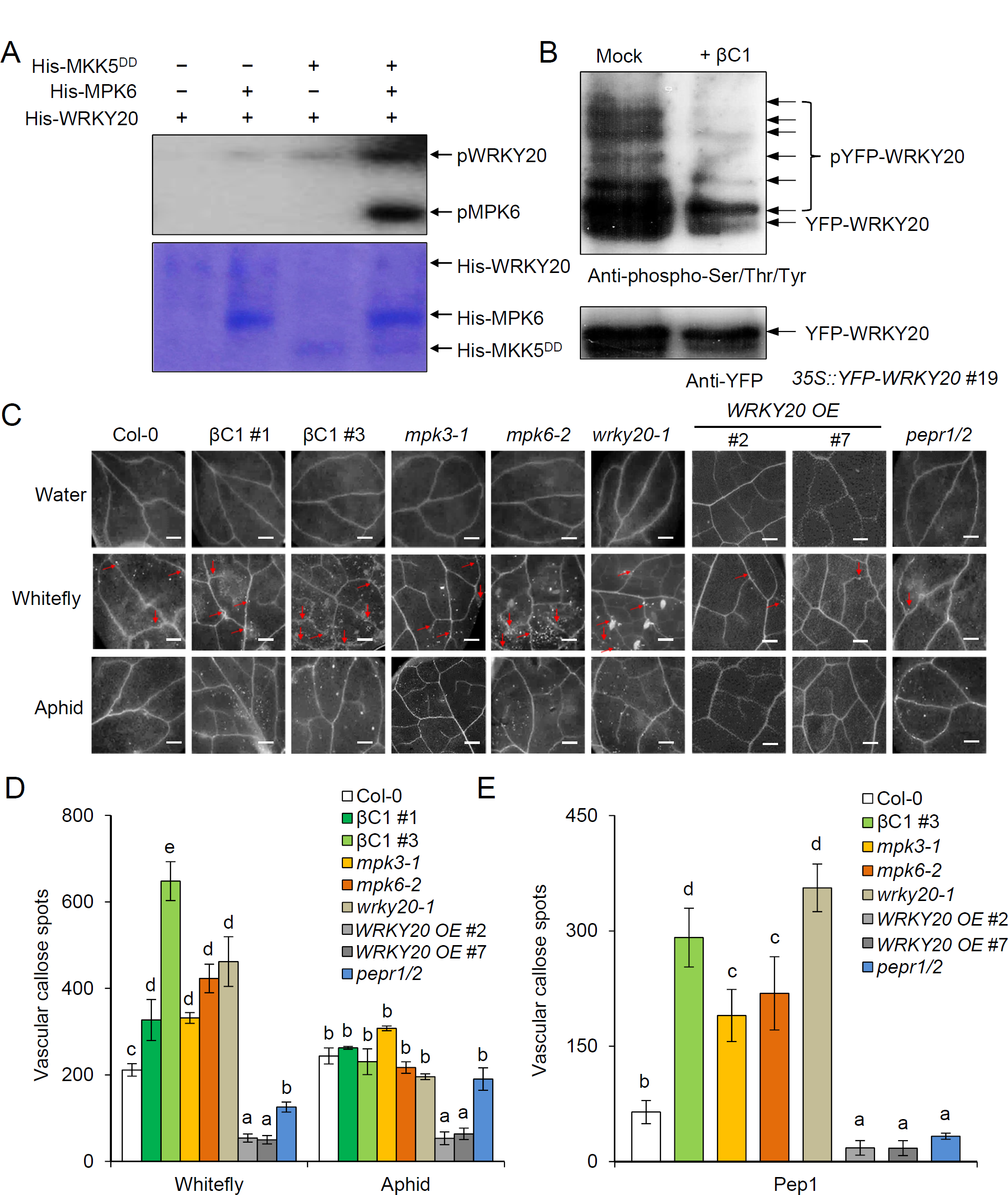
The MPK3/MPK6 substrate WRKY20 modulates callose deposition in a vascular-specific pattern. (**A**) *In vitro* phosphorylation of WRKY20 by MPK6. The phosphorylated form of WRKY20 (pWRKY20) was detected by in-gel kinase assay. The amount of input proteins was indicated by coomassie blue staining. (**B**) The phosphorylation levels of WRKY20 by treated with purified βC1 protein. The YFP-WRKY20 protein was extracted and purified from *35S:YFP-WRKY20* transgenic plants. The phosphorylation levels of WRKY20 were determined with anti-phos-Ser/Thr/Tyr antibody by immunoblot analysis. Loading amount of total YFP-WRKY20 protein was indicated by western blot using anti-YFP antibody. (**C**) Representative images of callose deposition in 10-day old *Arabidopsis* leaves upon water, whiteflies or aphids treatment for 12 hours. Scale bars = 300 μm. (**D**) The number of vascular callose spots by treated with water, whiteflies or aphids in *Arabidopsis* leaves was counted on each leaf using ImageJ software. Bars represent means ± SD (n=15). (**E**) The number of vascular callose spots by treated with Pep1 in *Arabidopsis* leaves. Bars represent means ± SD (n=15). Lowercase letters indicate significant differences among different lines according to one-way ANOVA followed by Duncan’s multiple range test (P < 0.05).

Serine or threonine followed by proline (SP or TP) is a minimal consensus motif for MAPK phosphorylation (Sharrocks et al., 2000). By bioinformatics analysis, we found ten potential MAPK phosphorylation sites in WRKY20 protein. To determine whether MPK3 or MPK6 phosphorylates WRKY20, we first conducted an *in vitro* phosphorylation assay. Figure 5A shows that WRKY20 can be phosphorylated by MPK6 *in vitro*, suggesting that WRKY20 is a direct substrate of MAPKs. We further conducted *in vivo* phosphorylation assay to check whether WRKY20 could be phosphorylated in plant cell. Total soluble WRKY20 proteins were purified by YFP-tagged beads. Multiple phosphorylation modification bands of WRKY20 could be more easily detected (Figure 5B). If βC1 could interfere with the interaction between MPK3 or MPK6 and WRKY20, then we would expect that phosphorylation cascade of MAPKs-WRKY20 will be weakened. Consistent with this prediction, phosphorylation levels of WRKY20 were obviously decreased when co-treated with purified βC1 protein in YFP-WRKY20 overexpression plants (Figure 5B). These results demonstrate that βC1 interacts with three host MAPK cascade proteins (MPK3/6 and WRKY20) and impairs this phosphorylation cascade from MPK3/6 to WRKY20.

### WRKY20 modulates callose response in a vascular-specific pattern

Upon pathogen or herbivore attacks, the defense phytohormone SA signaling is activated and callose deposition will then be enhanced to inhibit pathogen invasion (Fu and Dong, 2013). We have demonstrated that vascular-specific expressing WRKY20 could negatively regulate the SA biosynthesis and signaling pathway (Zhao et al., 2019). It would be expected that the hijacking of MPK3/6/WRKY20 by βC1 would activate plant SA signaling and enhances plant vascular callose deposition. Thus we performed a phenotypic assay for herbivore-triggered callose deposition in corresponding mutant plant lines and wild-type Col-0 *Arabidopsis* seedlings. Figure 5C,D and Figure 5-figure supplement 3A show that enhanced callose, in response to the infestation of whitefly in each of *mpk*3-1/mpk6-2/*wrky20-1* mutants as well as the βC1-expression plants, was spatiotemporally deposited in leaf vascular but not in mesophyll cell, while no obvious deposition occurred in the water-treated *Arabidopsis* seedlings. In contrast, no differences in vascular callose deposition were observed between the βC1-expression plants, *mpk*3/6/*wrky20* mutants, *pepr1/2* mutant and Col-0 plants upon green peach aphid infestation (Figure 5C,D). Consistently, a significant decrease in herbivory-induced callose deposition in the *WRKY20* overexpression lines compared with Col-0 plants, indicated that ectopic expression of *WRKY20* (*WRKY20 OE* #2 and #7) further suppresses the accumulation of vascular callose. We also observed reduction of vascular callose in the membrane receptor *PEPR1/PEPR2* mutant, suggesting that additional co-receptor was also involved in the sensing of whitefly herbivory.

Recently, pathogen-associated molecular patterns (PAMP) or damage-associated molecular patterns (DAMP)-induced callose deposition in cotyledons or leaves of *Arabidopsis* has emerged as a useful marker response to study plant defense response against herbivores and microbial pathogens (Clay et al., 2009; Luna et al., 2011). Therefore, we examined callose deposition in response to two different molecular patterns, bacterial PAMPs flagellin epitope Flg22 and a DAMP endogenous molecules Pep1. Consistent with some bacterial virulence effectors such as HopAI1 (Zhang et al., 2007), the viral βC1 also blocked flg22-induced callose deposition (Figure 5-figure supplement 3B,C). Further analysis revealed that Flg22- and Pep1-induced callose differ from that in βC1-plants. The viral βC1 did not affect Pep1-induced callose deposition. Interestingly, the pattern of Pep1-induced vascular callose response is similar to that of whitefly infestation in *Arabidopsis* mutants (Figure 5E and Figure 5-figure supplement 3C). The βC1-plants partially phenocopied two *mpk* mutants that increased Pep1-induced vascular callose deposition compared with Col-0 plants. The callose deposition in response to Pep1-treatment in mutant *wrky20-1* showed a vascular-specific pattern, further supporting its spatiotemporally repressive roles in callose deposition.

### The hijacking of WRKY20 by viral βC1 modifies whitefly feeding behavior and promotes virus transmission

Whitefly infestation increased callose accumulation at plasmodesmata, which in turn restricts the ability of whitefly to uptake nutrient from the plants (Fridborg et al., 2003; Zavaliev et al., 2011; Li et al., 2017). To respond to this host defense, herbivores secrete more saliva and therefore likely increase virus transmission. As shown in Figure 1E,F, the further activation of vascular callose response by begomoviral βC1 would positively affect whitefly feeding behavior and therefore enhance virus transmission. To further examine whether the enhanced vascular deposition upon whitefly feeding was related to virus transmission, we next performed electrical penetration graph (EPG) to track insect vector’s stylet activity and movement. The whitefly vector feeding behaviors mediate virus acquisition and inoculation by inserting its stylet bundle into a plant and locating along the stylets’ path towards the phloem (Prado Maluta et al., 2017). EPG recorded electrical resistance and electromotive force (Figure 6-figure supplement 1A). The EPG waveforms are engaged in intracellular punctures, xylem ingestion, salivation and phloem sap ingestion, as indicated by three distinct transmission cycle of vector-borne viruses including probing, virus inoculation and virus acquisition. Consistent with the activation of vascular callose response by βC1, the probing rate of whitefly on βC1-plants was higher than that on wild-type Col-0 plants, and that similar to the increased probing rate that was observed in *mpk3-1, mpk6-2* and *wrky20* mutants (Figure 6A and Figure 6-figure supplement 1B). Two phloem phases (salivation and phloem sap ingestion), referred to as E1 and E2 period waves, are associated with virus inoculation and acquisition by the insect vector, respectively. The total time for E1 and E2 periods by whiteflies was at least 2-fold higher in the *mpk* and *wrky20* mutant plants than those in Col-0 plants (Figure 6B,C). In addition, whiteflies exhibited longer periods of E1 and E2 periods on βC1-plants than those on the *mpk* mutant plants. The feeding behavioral changes observed on βC1-plants were similar to those described for tobacco (*Nicotiana tabacum*) plants infected with TA+β complex (He et al., 2015). These findings confirm that begomovirus βC1 manipulates whitefly feeding behavior on the host plant to facilitate virus transmission.

**Figure 6.**
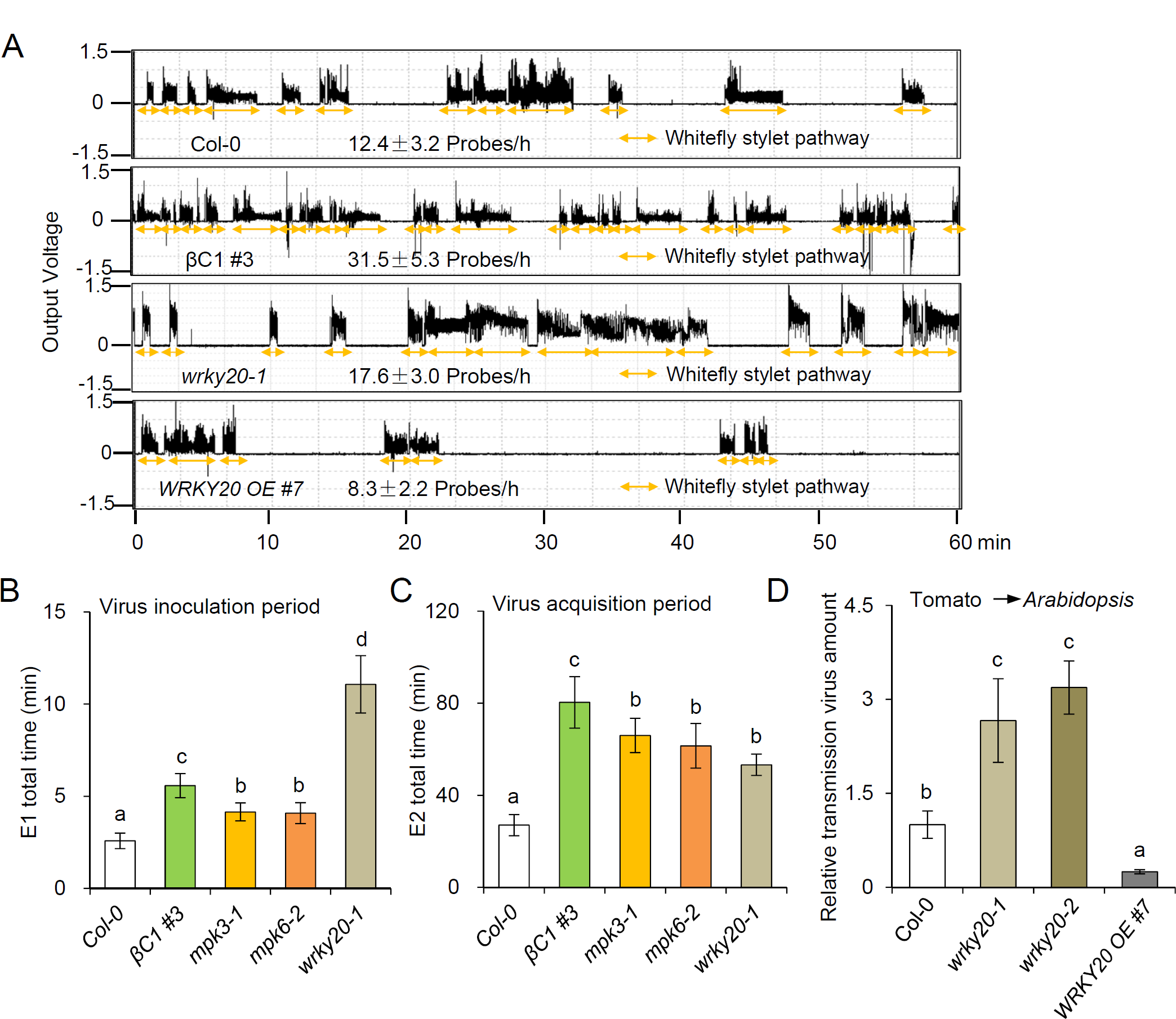
The hijacking of WRKY20 by viral βC1 modifies whitefly feeding behavior and promotes virus transmission. (**A**) Representative EPG waveforms of whitefly stylet pathway phases on *Arabidopsis* plants. Stylet pathways for whitefly represent stylet movement in the intercellular apoplastic space. Bars represent means ± SD (n=16). (**B, C**) The total E1 time (**B**) and total E2 time (**C**) of whitefly feeding on different genotypes of *Arabidopsis* plants was recorded for 8 h by electrical penetration graph (EPG) and analyzed by EPG software. Bars represent means ± SD (n=12). (**D**) The relative TYLCV virus amount transmitted from tomato to *Arabidopsis* plants by whiteflies. Bars represent means ± SD (n=8). Lowercase letters indicate significant differences among different lines according to one-way ANOVA followed by Duncan’s multiple range test (P < 0.05).

Mutant lacking *WRKY20* gene in *Arabidopsis* induced vascular callose response, we thus observed virus transmission among plants in *wrky20* mutants by viruliferous whiteflies. The quantity of virus transmitted into *wkry20* mutants was 3-4 fold higher than that in wild-type Col-0 plants (Figure 6D). *WRKY20* deficiency would make the host sensitive to virus dissemination by whiteflies. If this hypothesis is correct, then *WRKY20* overexpression should produce the opposite result. This was indeed the case in *Arabidopsis*. Overexpressing *WRKY20* in *Arabidopsis* plants showed reduced vascular callose deposition and reduced whitefly probing frequency (Figure 5C-E and Figure 6A). The quantity of virus in *WRKY20* overexpression plants was reduced to only 25% of that in Col-0 plants (<u>Figure 6D</u>). Together, the results demonstrate that βC1-WRKY20 interaction specifically promotes begomovirus transmissibility and virulence by modulating the tissue-specific defense in *Arabidopsis*.

## Discussion

Although plants are often in continual contact with potential herbivores and pathogens, a successful infection is rare. However within the last a few decades, some begomoviruses have emerged as important agents of plant virus diseases worldwide (Zhou, 2013; Yang et al., 2019). In this study we have characterized an important plant MPK-WRKY20 phosphorylation cascade against begomoviruses and the whitefly vector. Although our conclusion mainly based on the model plant *Arabidopsis*, factors of this signaling are conserved at least in higher core eudicots plants. When whitefly or begomovirus infects host plants, plant rapidly triggers some early defense signals, including kinases activation such as MAPK and SnRK, Ca^2+^ elevation and ROS burst. Active MAPKs interact with and phosphorylates WRKY20. WRKY20 is a vascular-specific transcription factor that physically and functionally interacts with ORA59, a key intersection point of pathways involving defense phytohormones, i.e., JA, ethylene, and SA (Zhao et al., 2019). The deficiency of *wrky20* or transgenic *βC1* expression increases expression of and SA-responsive genes (*AtPAL1, AtSID2, AtPAD4, AtEDS5*, and *AtPR1*) and SA synthesis. The repressed SA signaling by WRKY20 leads to suppression of callose response. Possible repressive roles in callose deposition confer changes of whitefly feeding behavior, and therefore affect virus inoculation and acquisition (Supplementary Figure 4A,B). Furthermore, βC1 is phosphorylated by plant SnRKs mainly on serine^33^, resulting in a low phosphatase activity. The βC1^S33D^ still can interact with βC1 substrates (MAPKs and WRKY20) (Figure 2-figure supplement 3C and Supplementary Figure 3).

Interestingly, when a begomovirus infects host plant, the viral βC1 attracts whiteflies by hijacking a key component in the JA pathway MYC2-mediated olfactory defense against herbivores (Luan et al., 2013; Li et al., 2014). Plant responses induced by herbivory, e.g. whitefly feeding, is important for virus to suppress host defense. Once whitefly feeding perceived by the plant receptors PEPR1/2, the begomovirus-induced transcription of plant *SnRK* genes is suppressed by whitefly herbivory. This suppression of expression of host *SnRK* genes alleviates the phosphorylation of βC1 at serine^33^. Alternatively, the co-infection by whitefly and begomovirus may activate some uncharacterized host phosphatase of βC1 that dephosphorylates it and converts it into transmission state with higher phosphatase activity. Begomovirus-encoded βC1 is rapidly activated by whitefly infestation, and impairs the phosphorylation cascade of MPK3/6-WRKY20. Accordingly, the early plant defense responses, e.g. MAPK activation, Ca^2+^ elevation and ROS burst, are suppressed, thereby promoting begomovirus accumulation. SA-regulated vascular callose response is activated by βC1, thus altering whitefly feeding behavior. Finally, βC1 confers easier whitefly-mediated virus transmission between plants by HIDS (Supplementary Figure 4C).

Post-translation modification (PTM) has been widely proven to be a powerful and rapid antiviral mechanism both in plants and human (Liu et al., 2016). Interaction of a viral product with a host protein involved in the defense response results in changes in phosphorylation or ubiquitination, cleavage or degradation, but these specific biochemical mechanisms for inhibitory effects are still illusive. In this study, we characterized the begomovirus pathogenicity factor βC1 as the first phosphatase encoded by plant viruses. To date, more than 1300 full-length β-satellite sequences, belonging to 61 species of the genus *Betasatellite*, have been characterized from various host plants in 37 countries (Yang et al., 2019). The βC1 protein can reprogram cellular processes in the host plant and facilitate begomovirus infection and transmission, mainly by interacting with multiple host proteins. Our results here demonstrated three substrates of βC1 phosphatase, two kinases MPK3/6 and one transcription factor WRKY20. Both MPK3/6 and SnRK kinases phosphorylate MYC2 (Sethi et al., 2014; Wurzinger et al., 2018), another interactor of βC1. Phosphorylation of MYC2 is essential for its function to regulate gene transcription. Additional to known ETHYLENE RESPONSE FACTOR6 (ERF6) (Xu et al., 2016), WRKY20 as another transcription factor substrate for MPK6 to regulate the biosynthesis of metabolite glucosinolates, is important for the plant defense response against herbivores and microbial pathogens. The hijacking of plant transcription factor network by the βC1 phosphatase reshapes the ecological community of plants via changing plant chemicals, such as glucosinolates, volatile organic compounds (e.g., terpenoids), phytohormones (e.g., ethylene), and toxic polypeptides (e.g., defensin and pathogenesis-related protein 1).

Based on the findings of the function of βC1 in tripartite interactions reported in this study, it will be interesting to characterize new substrates of βC1 to fully understand how significant of the manipulated phosphorylation network by this pathogenicity factor during begomovirus infection and disease epidemic, as well as to reexamine known interactors of βC1. It seems like that the phosphatase activity of βC1 might be involved in its roles in host autophagy process, but might not be associated with its known silenceing suppressor activity. Here we show that phosphatase activity of βC1 protein looks unrelated with its known silencing suppressor activity (Figure 2 and Supplementary Figure 1). Meanwhile a mutation of the neighbor amino acid valine^32^ (next to the active site serine^33^) of βC1^V32A^ abolished its interaction with a key autophagy protein NbATG8f, and begomovirus carrying βC1^V32A^ showed increased symptoms and viral DNA accumulation in plants (Haxim et al., 2017). Nevertheless, further work is required to further distinguish the relationship between phosphorylation and autophagy or RNA silencing machineries in controlling begomovirus pathogenesis. In addition, proteomic analysis in combination with metabolic analysis of βC1-expressing plants, *myc2-1, wrky20*, and mutations in autophagy machinery will also be very helpful to ascertain the exact roles of autophagy, MYC2 and WRKY20 in βC1-mediated whitefly-begomovirus mutualism.

The significance of this HIDS in nature and agricultural ecology warrants more attention. The effect on pathogen transmission could be extended beyond begomoviruses to broader fields of vector-borne parasites. It may also explain a common phenomenon, Viral Perceptive Behavior (VPB) by which a virus senses and responds to the presence of a vector identified in other arboviruses (Martiniere et al., 2013; Bak et al., 2017). Moreover, our research presents fascinating biochemistry opportunities to explore the roles of post-transcriptional modification (PTM) on pathogen proteins in response to the presence of a vector. PTM, especially phosphorylation regulates the “arms race” between viruses and plants/animals (Liu et al., 2016; García-Sastre, 2017). We have shown that plant begomoviruses enhance its own transmission in a dephosphorylation-dependent HIDS manner. It seems that a switchable phosphorylation on serine^33^ of βC1 is crucial for the suppression of the MAP kinase-mediated plant defenses and confers adaptation advantages to begomoviruses. Therefore, we can explore the adaptive significance of virus effects on hosts and vectors by evaluating virus effects across phylogenetically diverse viruses that share a transmission mechanism. It may provide predication on whether and how one kind of invasive arbovirus can be prevail under certain environmental condition. Frequent high nucleotide substitution results in high transmission efficiency in virus evolution, such as mosquito-transmitted ZIKA virus (Liu et al., 2017). These studies will enrich our understanding of the evolution of parasite manipulation and facilitate development of new epidemiological models for predicting and mitigating virus threats to human and agriculture.

In summary, our study identified the MPK-WRKY20 phosphorylation cascade as one of host arsenals against whiteflies and begomoviruses. The data here are limited to laboratory observations; nevertheless, they provide evidence for how this HIDS could take place to facilitate begomovirus rapid dissemination. This type of tripartite interactions examined in this study likely occurs widely in natural and agricultural ecosystems, our findings provide new insight into the tripartite interactions between parasite, vector and host, and offer clues for establishing biotechnological ways to the development of antivirals for containing animal-vectored pathogens.

## Materials and Methods

### Plant materials and growth condition

*Arabidopsis thaliana* ecotype Columbia (Col-0, WT) was used in this report. The transgenically expressing *βC1* plants (βC1 #1 and βC1 #3) (Yang et al., 2008), *mpk3-1* (Zhang et al., 2007), *mpk6-2* (Liu and Zhang, 2004), *pepr1/2* (Liu et al., 2013), *wrky20-1, wrky20-2, WRKY20* overexpression lines driven by its native promoter (*WRKY20 OE#2, #7*) lines (Zhao et al., 2019) and *35S:GCaMP3* (Col-0/*GCaMP3*) mutants (Vincent et al., 2017) have been previously described. The βC1 #3*/GCaMP3, mpk3-1/GCaMP3, mpk6-2/GCaMP3* or *pepr1/2/GCaMP3* mutants were generated by crossing *35S:GCaMP3* line with the corresponding mutant background, which confirmed with genotyping. Homozygous lines were used in all these experiments. *Arabidopsis* plants were grown in a growth chamber at 22°C with a 12 h light/12 h darkness cycle. *N. benthamiana, S. lycopersicum* (Zhongza no.9; tomato) and *Gossypium barbadense* (Xinhai no. 21; cotton) plants were grown in a growth chamber at 25°C with a 12 h light/ 12 h darkness cycle.

### Virus inoculation

For begomovirus infection experiments, *N. benthamiana*, cotton or *Arabidopsis* plants were inoculated with *Agrobacterium* EHA105 carrying CA+Cβ (Jia et al., 2016), TA+β (Li et al., 2014), with two βC1 mutant β-satellites (TA+mβ and TA+β^S33D^). For begomoviral pathogenesis, viral DNA was detected by PCR with TYLCCNV-specific primers (Supplementary Table 1) to identify the infected *Arabidopsis* plants at 21 days post TA+β infection.

For Tobacco rattle virus infection, leaves of 3-week-old *N. benthamiana* plants were agroinfiltrated with psTRV1 and psTRV2 (Qu et al., 2012).

### Virus transmission assays

A group of 30 viruliferous adult whiteflies from TYLCV-infected tomato plants were captured and placed in a covered cup on plant. After 72 h feeding on transmission target plants, viruliferous whiteflies were then gently removed off. Whole plant was collected for further virus analysis. Viral DNA was quantitated through real-time PCR with TYLCV-specific primers (Supplementary Table 1).

For virus infection efficiency, a group of 30 TYLCV-viruliferous whiteflies were captured and fed on *Arabidopsis* for 72 h challenge. Plant infected ratio were calculated by plants with PCR positive with TYLCV-specific primers against the total challenged plant.

### Insect infestation

The population of MEAM1 (Middle East-Asia Minor 1) whiteflies were maintained on uninfected tomato plants in the lab as reported before (Li et al., 2014). For whitefly infestation, 50 adult whiteflies were placed in a leaf-clip cage on an undamaged plant leaf for an indicated treatment time.

For aphid infestation, six-week old *Arabidopsis* plants were placed undamaged in a leaf-clip cage. Thirty adult green peach aphids (*Myzus persicae*) were then placed into each leaf cage for infestation treatment. The whole treated plant was gently collected and kept in liquid nitrogen for further analysis.

### Quantitative RT-PCR

Total RNA was isolated from plants using the RNeasy plant mini kit (Qiagen, 74904) including on-column DNase treatment. Reverse transcription was performed using 2μg RNA of each sample and oligo (dT)_18_ primers by M-MLV reverse transcriptase (Promega, 28025013). Three independent biological samples were collected and analyzed. Real-time PCR was performed on a Bio-Rad CFX96 real-time PCR system with Thunderbird™SYBR qPCR mix (TOYOBO, QPS-201) and gene-specific primers listed in Supplementary Table 1. The *Arabidopsis ACTIN2* and *N. benthamiana NbEF1α* were used as internal control.

### Plant protein extraction and in-gel kinase activity assay

Plant soluble protein was extracted from leaf samples using extraction buffer (20mM Tris-HCl pH8.0, 100mM NaCl, 10mM MgCl_2_, 50 mM DTT, 0.5mM PMSF and protease inhibitor cocktail). Equal amount of total protein was loaded and separated on 10% SDS-polyacrylamide gels and transferred to a polyvinylidene difluoride (PVDF) membranes (Millipore, IPVH00010). MAP kinase activation was detected by western-blot analysis with anti-phospho-p44/42 MAPK (Erk1/2) (Thr202/Tyr204) antibody (Cell Signaling Technology, 9101S) and anti-rabbit horseradish peroxidase-linked sheep antibody. Immunoblot signals were detected by Amersham ECL prime kit (GE Healthcare, RPN2232). Relative activation of MAPKs was analyzed by Image J software from three independent experiments.

### Recombinant protein purification

DNA fragments encoding full-length of MPK3, MPK6, WRKY20 and βC1 were amplified by PCR with Phusion high-fidelity DNA polymerase (Thermo Scientific, F530S) using primers listed in Supplementary Table 1. These amplified DNA fragment was ligated into pGEX-DC or pET-28a (+) to generate GST fusion or His fusion constructs. Point mutation in specific amino acids of βC1^S33D^ was generated by Site-directed mutagenesis kit (Sangon Biotech, Shanghai) and His-MKK5^DD^ (T215D/S221D) construct was obtained from Dr. Jie Zhang (Zhang et al., 2007). Recombinant proteins were produced and purified by method used before (Zhao et al., 2019), according to the manufacturer’s instructions.

### *In vitro* kinase assay

The *in vitro* kinase assay was performed as previously described (Zhang et al., 2007). Recombinant His-MPK6 protein was incubated with or without a constitutively active His-MKK5^DD^ at 30°C for 30 min in a kinase reaction buffer (20 mM Tris -HCl pH 7.5, 1 mM DTT, 10 mM MgCl_2_, 1 mM CaCl_2_, 50 μM ATP). The phosphorylated His-MPK6 protein was then incubated with different amounts of purified His-βC1 protein or His-WRKY20 protein in the kinase reaction buffer with [ϒ-32P]-ATP (1μCi per reaction) at 30°C for 30 min. The reactions were stopped by the addition of 4×NuPAGE LDS sample buffer. Phosphorylated His-MPK6 proteins or His-WRKY20 proteins were visualized by autoradiography in a 10% SDS-PAGE gel.

### *In vitro* pull-down assay

The *in vitro* binding assay was performed as preciously described (Ye et al., 2015). Pull-down proteins were separated on 10 % SDS-polyacrylamide gels and detected through immunoblotting with anti-GST antibody (TransGen Biotech, HT601-02).

### Phosphatase activity assays

Fifty micromolar synthetic MAPK phosphopeptide (SESDFM-pTE-pYVVTR) (SciLight Biotechnology, Beijing) was incubated with 25 μM recombinant His-βC1 or His-βC1^S33D^ protein in a 50 μl reaction buffer containing 10 mM HEPES (pH 7.4), 150 mM NaCl, and 1 mM EDTA and incubated at 30 °C for 30 min. Mass of βC1-treated phosphopeptide was analyzed using matrix-assisted laser desorption/ionization time-of-flight mass spectrometry. The specific modification of phosphor-Thr and phosphor-Tyr residues in the treated peptide was measured by tandem mass spectrometry (Thermo Scientific™ Orbitrap Fusion™, QLbio company, China), and the intensity of phosphopeptide was further analyzed by MaxQuant software.

### Phos-tag SDS/PAGE and immunoblot analysis

Leaves of six-week-old *N. benthamiana* plants were agroinfiltrated by *A. tumefaciens* cells harboring *myc-βC1-myc* or *myc-βC1*^*S33D*^, and incubated for 72 hours in a growth chamber at 25°C with a 12 h light/ 12 h darkness cycle. Plant samples were collected and ground into powder in liquid nitrogen, following with adding 2×NuPAGE LDS sample buffer (Invitrogen, NP0008) containing 0.05mL/mL β-mercaptoethanol, and protease and phosphatase inhibitor cocktail (Pierce, A32959) on ice for 10 min. The suspension containing total myc-βC1 or myc-βC1^S33D^ protein was boiled at 100°C for 5 min, and then centrifuged at 14,000g at room temperature for 5min.

For Phos-tag gel, the special 12% SDS-PAGE gel was prepared containing 100 μM ZnCl_2_ and 50 μM Phos-tag acrylamide AAL-107 (WAKO, 304-93521) according to the manufacturer’s instructions. Western blot analysis with Phos-tag gels was performed as previously reported with some modifications (Kinoshita et al., 2006). Put the gel running apparatus in ice baths and the protein gel was first run at 50V for 6 hours, then incubated in transfer buffer (48 mM Tris-base, 39mM Glycine, 20 % MeOH) containing 10 mM EDTA for 10 min for three times with gently shaking. The gel was next incubated with gentle agitation in transfer buffer containing 0.05% SDS for another 10 min. Then the gel was electrophoretic transferred onto PVDF membrane using a wet-tank blotting apparatus at 100 V for 3 hours. The immunoblot was probed with anti-myc antibody.

### Yeast two-hybrid analysis

Full-length protein of βC1 or WRKY20 was cloned into the pGBT9 vector through LR reaction to generate BD-βC1 or BD-WRKY20. Full-length protein of *Arabidopsis* MPK3/MPK6 was cloned into the pGAD424 vector through LR reaction to generate AD-MPK3/6. The yeast strain Y2H Gold was co-transformed with BD-βC1 or BD-WRKY20 and AD-MPK3/6 constructs and plated on SD/-Leu/-Trp selective dropout medium. Colonies were transferred onto SD/-Leu/-Trp/-His with 20 mM 3-amino-1,2,4,-triazole (3-AT) plates to verify positive clones. The empty vectors pGBT9 and pGAD424 were used as negative controls.

### Bimolecular fluorescence complementation (BiFC) assay

The DNA fragments of MPK3, MPK6, WRKY20 and βC1 or βC1^S33D^ were cloned into pGreen-pSAT1 serial vectors to generate fusion genes with nEYFP or cEYFP at the amino- or carboxy-terminus with a cauliflower mosaic virus *35S* promoter. All constructs were transferred into *Agrobacterium* C58C1 competent cells. The BiFC assay was performed as previously described (Zhao et al., 2019). *Agrobacterial* cells containing indicated constructs were infiltrated into leaves of three-week old *N. benthamiana*. Images of fluorescence were taken using a confocal microscopy (Leica SP8) after 48 h incubation.

### Co-immunoprecipitation (Co-IP) assay

Constructs containing βC1-myc, YFP, YFP-MPK3, or YFP-MPK6 were infiltrated with *agrobacterium* strains (EHA105) and transiently expressed in leaves of four-week-old *N. benthamiana*. Total proteins were extracted from infiltrated leaf patches in 1 ml lysis buffer [50mM Tris-HCl pH7.4, 150mM NaCl, 2mM MgCl_2_, 10% glycerol, 0.5% NP-40, 1mM DTT, protease inhibitor cocktail (Roche, 04693159001)] after 48 h co-expression. Twenty milligram protein extracts were taken as input and the rest of the extracts were incubated with the GFP-Trap beads (ChromoTek, gta-20) for 1.5 h at 4°C. Immunoblot analysis was performed with anti-myc and anti-YFP antibodies.

### Active oxygen species ROS

ROS released by leaf tissue was assayed as described (Gomez-Gomez et al., 1999). Seven-week old *Arabidopsis* leaves were sliced into approximately 3 mm leaf discs, incubated in 200 μl water in a white 96-well plate. After incubation for 12 hours, One hundred reaction solution with 1 μM Pep1, 100 μM luminol and 10 μg ml^-1^ horseradish peroxidase (Sigma) plus was added to each well. Luminescence was recorded immediately for 30 min by Microplate Luminometer (Promega).

### H_2_O_2_ detection with DAB staining

Seven-week old *Arabidopsis* leaves with or without herbivore infestation for 6 h were stained with 1 mg/ml DAB-HCl solution (Sigma, D5637) overnight in darkness. The samples were then destained in the ethanol until leaves turned clear. Images were then captured with a stereoscopic microscope.

### Ca^2+^ signal analysis

Leaves of three-week old plants were detached using sharp scissors and placed in wells of a black 96-well assay plate (Corning costar, 3916) with 200 μL of distilled water. Abaxial surface faced up. Plates were then kept at room temperature 3 hours to dissipate wound-induced Ca^2+^ signals. A final concentration of 1 μM Pep1 was used to induce Ca^2+^ level in each plant. A multiscan spectrum (480-525nm) was used to quantify fluorescence strength which indicates cytoplasmic free calcium (Ca^2+^) signal upon induction in each *35S:GCaMP3* expressed plants. The Ca^2+^ level was represented as changed fluorescence ratio at each time point (ΔF/F), which was calculated according to the equation ΔF/F = (F-F0)/F0, where F0 was the average baseline fluorescence calculated from the average of F over the first 30 frames of the recording (Vincent et al., 2017).

### Callose staining

Seven-week old *Arabidopsis* leaves were infiltrated with water, 1 μM Pep1, 1 μM Flg22, or ten-day old *Arabidopsis* cotyledon were treated by whiteflies and aphids. Leaves were harvested after treatment for 12 hours, cleared, and stained with aniline blue for callose as previously described (Adam and Somerville, 1996). Leaves were mounted in 50% glycerol, and epifluorescence was visualized with a fluorescence microscope under ultraviolet light. The number of callose deposits was counted by Image J software.

### Electrical penetration graph (EPG)

Adult female whitefly was attached to the Giga-8 EPG system (EPG-Systems, Wageningen University, Netherlands) using 12.7 μm diameter of gold wire and silver glue. The whitefly was placed on six-week old *Arabidopsis* plants. One whitefly was added to each plant, and this represented one biological replicate. Feeding behavior of whitefly on *Arabidopsis* was recorded for 8 h using Stylet+d (EPG Systems) and data were analyzed with the software Stylet+a (EPG Systems). Relevant EPG parameters were calculated using the Microsoft Excel spreadsheet.

## Data Analysis

Differences in genes expression level, relative MAPK activity and relative fluorescence intensity were analyzed using a two-tailed Student’s *t*-tests for comparing two treatments or two lines. Significant differences among different lines in number of callose spots, vascular callose spots, normalized GFP fluorescence (ΔF/F), relative luminescence (RLU), whitefly feeding behaviors, relative transmission virus amount, transmission efficiency and relative virus titer were tested using one-way ANOVA following by Duncan’s multiple range tests.

## Acknowledgements

We thank Prof. Jinlong Qiu [Institute of Microbiology, Chinese Academy of Sciences (IMCAS)] for *mpk3-1* and *mpk6-2* mutants, Prof. Jian-Min Zhou (Institute of Genetics and Developmental Biology, Chinese Academy of Sciences) for *pepr1/2* mutant, Prof. Daniel J. Kliebenstein (University of California, Davis, USA) for suggestions, Profs. Jie Zhang and Jun Liu (IMCAS) for invaluable assistance with experiments and suggestions.

## Additional information Competing interests

The authors declare no competing interests.

## Author contributions

Jian Ye, Pingzhi Zhao and Ning Wang conceived and designed the project. Pingzhi Zhao and Ning Wang conducted most of experiments. Insect assays were performed with the help of Xiangmei Yao. Saskia A. Hogenhout provided the *35S:GCaMP3* parental line used for crossing starting materials. All authors analyzed the data and reviewed the manuscript. The manuscript was written by Jian Ye, Pingzhi Zhao and Ning Wang with contributions from Changxiang Zhu, Xueping Zhou, Shu-sheng Liu, and Rongxiang Fang.

## Additional files Supplementary files

**Supplementary Figure 1.**
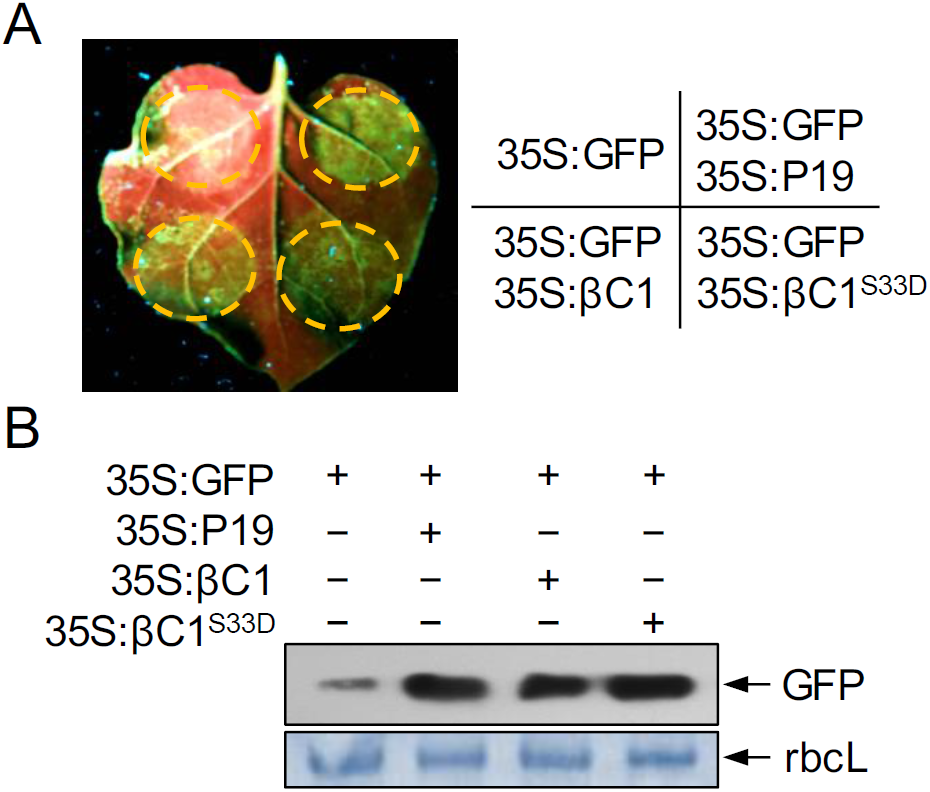
A phosphorylation mimic βC1^S33D^ does not affect the RNA silencing suppressor activity of βC1. (**A**) RNA silencing suppressor activity assay in *N. benthamiana* plants. Leaves were co-agroinfiltrated with constructs harboring GFP (*35S:GFP*) plus either 35S: TBSV P19 (*35S:P19*), βC1 (*35S:βC1*), or βC1^S33D^ (*35S:βC1*^*S33D*^). TBSV, Tomato bushy stunt virus. The agroinfiltrated leaves were photographed under UV light at 3 dpi. (**B**) Western blot assay of GFP accumulation in agroinfiltrated leaf patches shown in panel a. Stained membrane bands of the large subunit of Rubisco (rbcL) were used as a loading control.

**Supplementary Figure 2.**
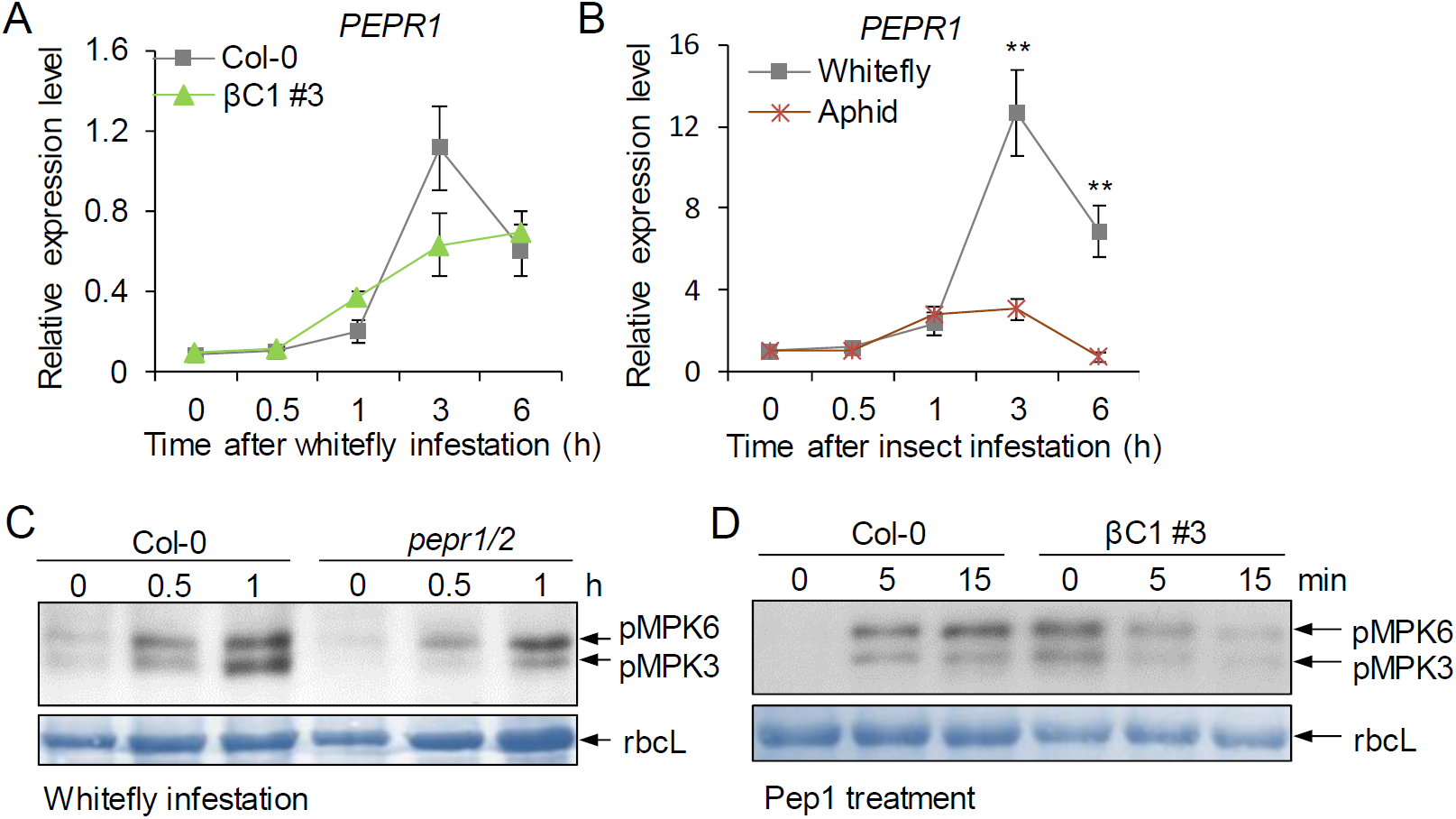
The receptors PEPR1 and PEPR2 involve in the perception of whitefly-triggered immunity. (**A**) Relative expression level of *PEPR1* upon whitefly infestation in Col-0 and βC1 #3 plants. Bars represent means ± SD (n=4). (**B**) Relative expression level of *PEPR1* upon whitefly infestation or aphid infestation in Col-0 plants. Bars represent means ± SD (n=4) (**, P< 0.01; Student’s *t*-tests). (**C**) Whitefly-induced MAPK activation is reduced in *pepr1/2* mutants. Three-week-old plants were infested with whiteflies for the indicated periods. The activities of *Arabidopsis* MPK3 and MPK6 were determined by immunoblot analysis using anti-phospho-p44/42 antibody. (**D**) βC1-induced MAPK activation is suppressed by Pep1. *Arabidopsis* plants were treated with 0.1 μM Pep1 for the indicated periods and immunoblotting was performed using anti-phospho-p44/42 antibody.

**Supplementary Figure 3.**
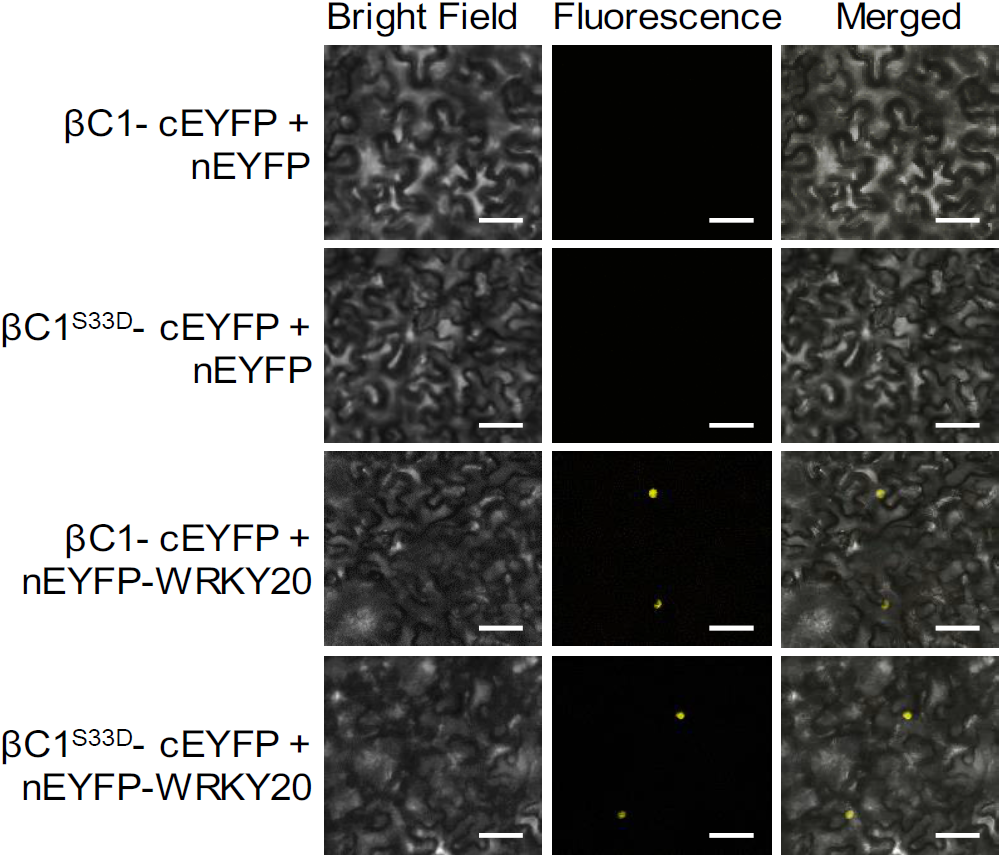
The phosphorylation mimic mutant βC1^S33D^ does not influence the interaction with WRKY20. BiFC analysis shows interaction between wild-type βC1 and WRKY20 or the phosphorylation mimic mutant βC1^S33D^ and WRKY20. Fluorescence was observed when βC1 or βC1^S33D^ fused with the C-terminal part of EYFP complemented WRKY20 fused with the N-terminal part of EYFP. Unfused nEYFP was used as a negative control. Scale bars = 50 μm.

**Supplementary Figure 4.**
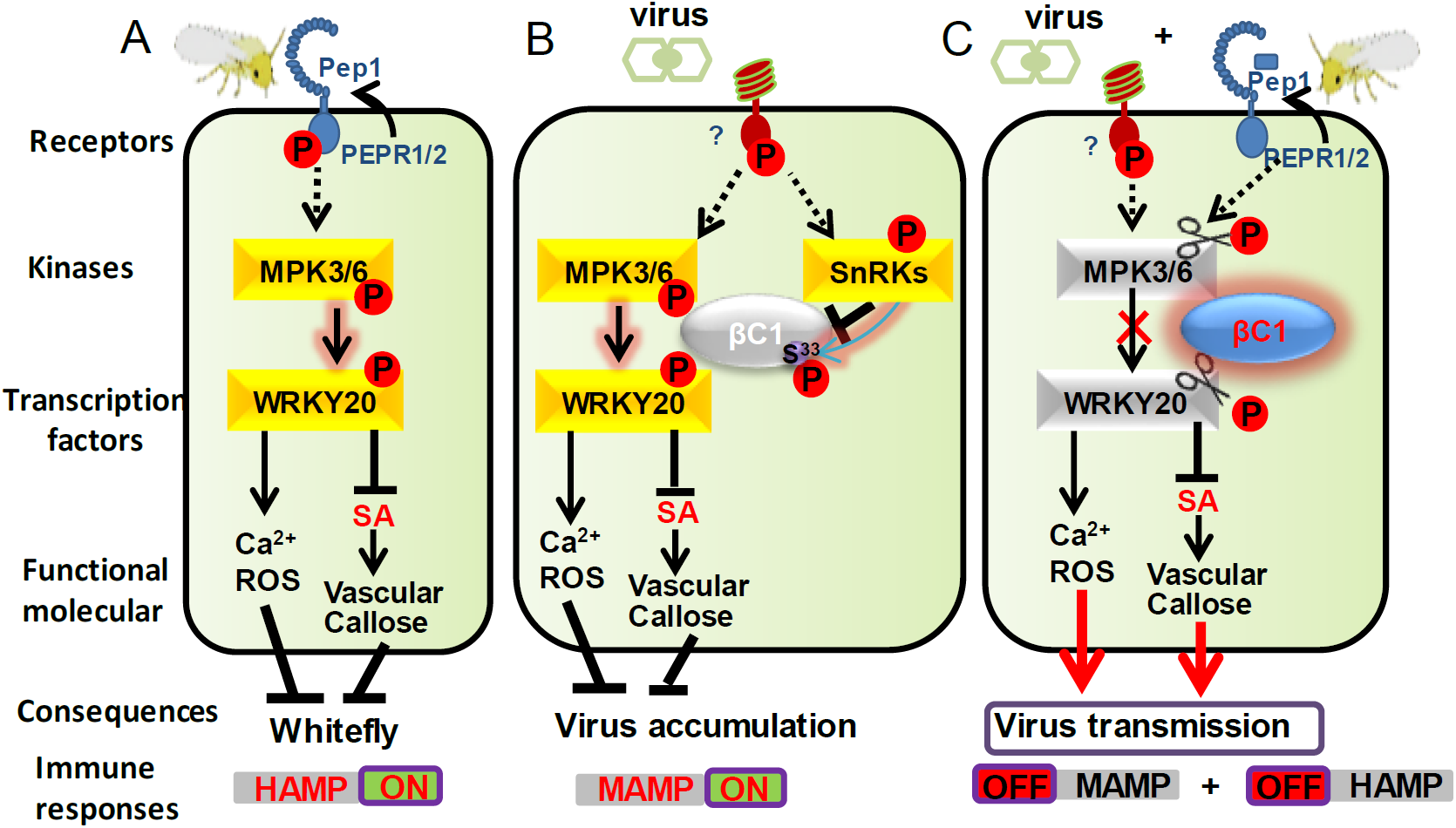
A working model for herbivore-induced viral phosphatase disarming plant antiviral immunities to facilitate pathogen transmission. (**A, B**) When whitefly (**A**) or begomovirus (**B**) infects host plants, plant rapidly triggers some early defense responses, including MAPK activation, calcium elevation and ROS burst, thus restricting virus accumulation in the host plant. Active MAPKs interacts with and phosphorylates downstream transcription factor WRKY20. WRKY20 is a vascular-specific transcription factor and suppresses SA signaling which leads to decreases vascular callose deposition, thus confers changes of whitefly feeding behavior and limits virus inoculation and acquisition. Meanwhile, betasatellite-encoded βC1 is at a low phosphatase activity phosphorylated by plant SnRKs. (**C**) Once whitefly infestation perceived by plant receptors PEPR1/2 in virus-infected host plant, the phosphatase activity of βC1 is rapidly activated by whitefly infestation, and impairs the phosphorylation cascade of MPK3/6-WRKY20. Accordingly, the early plant defense responses, e.g. MAPK activation, Ca^2+^ elevation and ROS burst, are suppressed, thereby promoting begomovirus accumulation. SA-regulated vascular callose response is activated by βC1, thus altering whitefly feeding behavior. Finally, βC1 confers easier whitefly-mediated virus transmission between plants by herbivore-induced host defense suppression.

**Supplementary Table 1.**
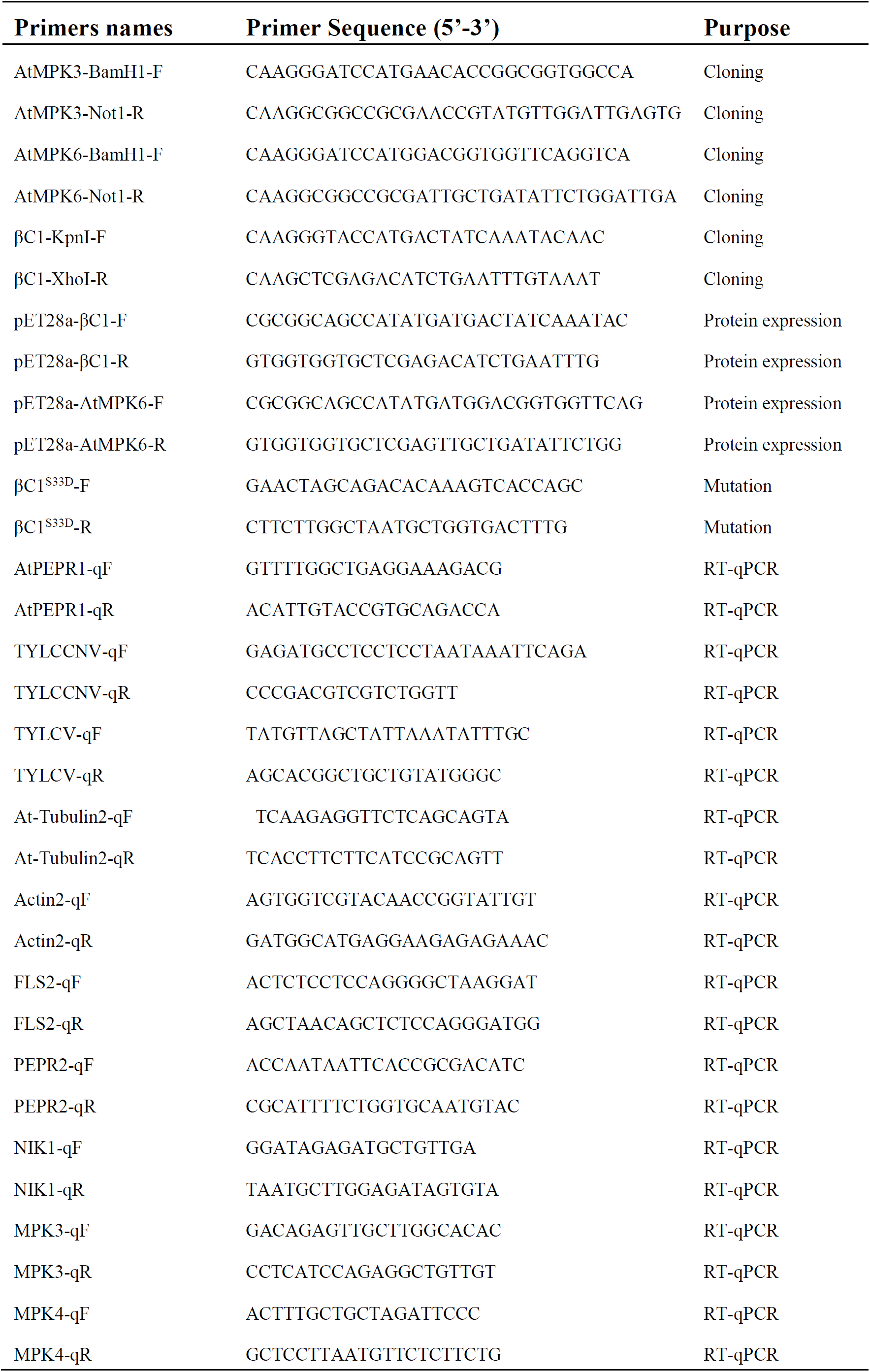
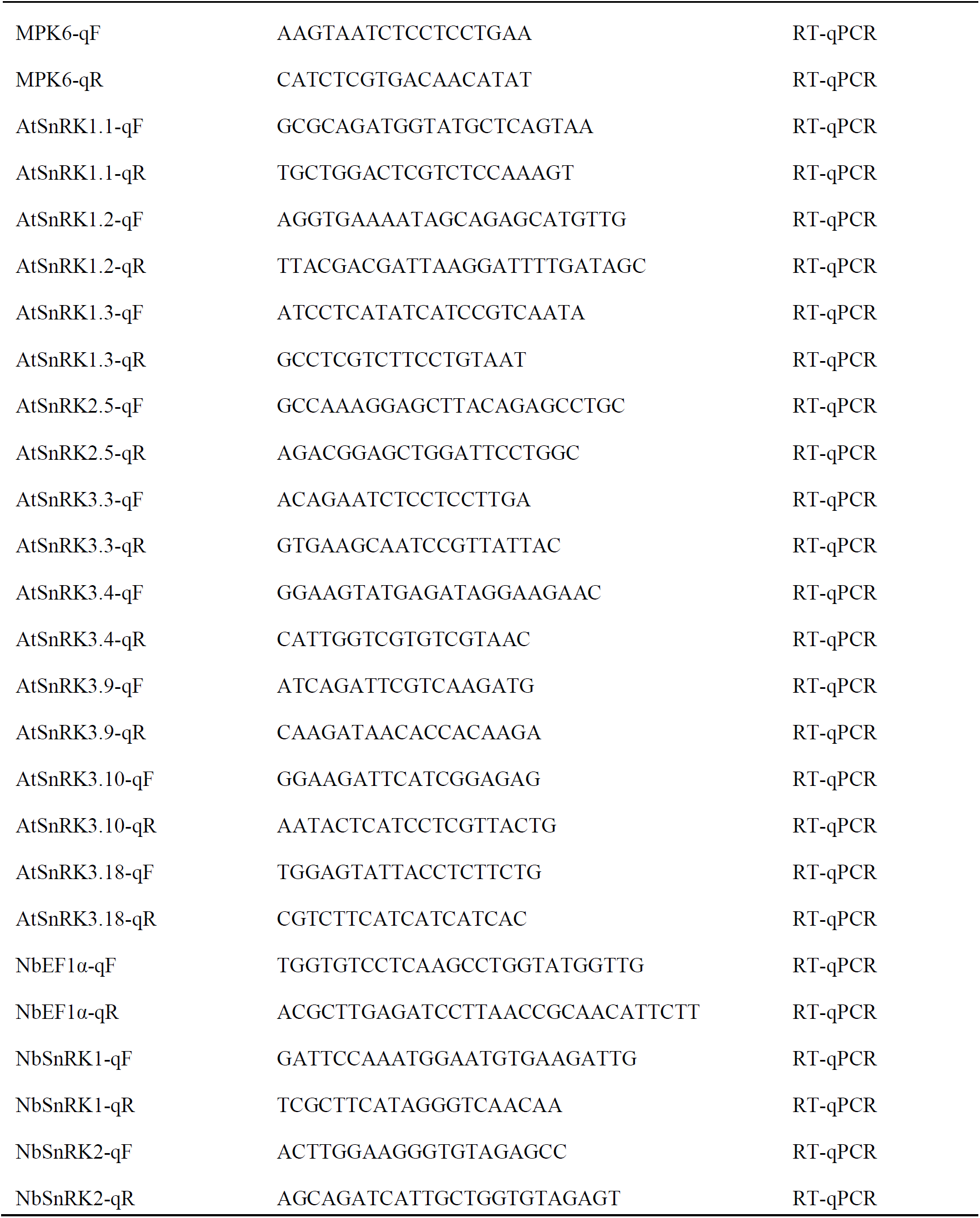
Primers sequences used in this investigation.

## Figure legends

**Figure 1-figure supplement 1.**
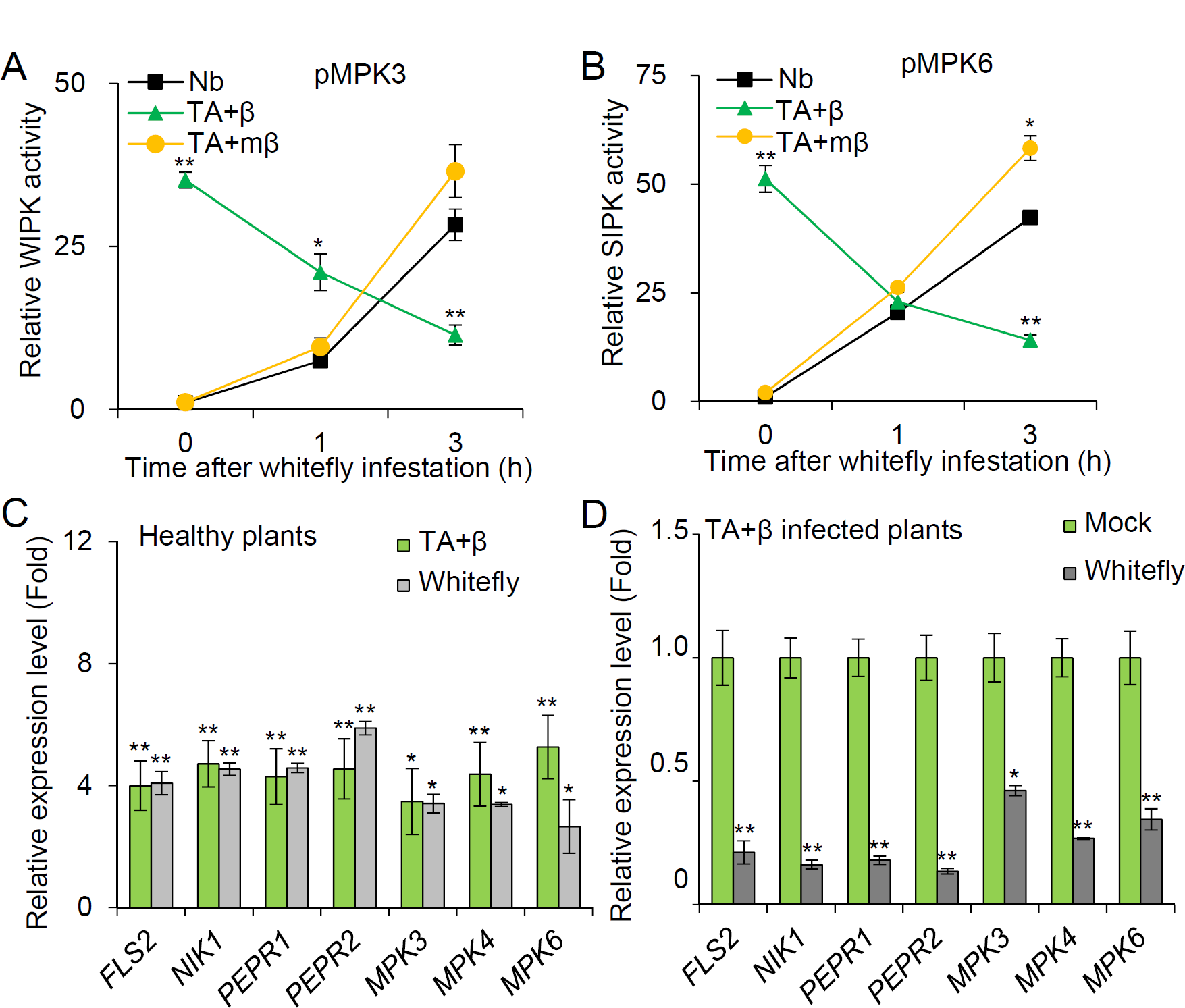
Whitefly herbivory induces defense suppression in begomovirus-infected plants. (**A, B**) Quantitative analysis on activities of MPK3 (**A**) and MPK6 (**B**) upon whitefly infestation in healthy *N. benthamiana* (Nb) plants or Nb plants infected by TYLCCNV complex (TA+β) or TYLCCNV plus βC1 betasatellite mutant complex (TA+mβ). MAPK activity was analysied by ImageJ software to quantify band intensities from the immunoblot analysis in Figure 1B. (**C**) The expression of genes associated with plant immune responses in *Arabidopsis* upon TA+β infection or whitefly infestation. Healthy Col-0 plants were inoculated with TA+β at 14 days post inoculation (dpi) or infested with whitefly for 6 h before sampling. Relative expression levels (fold changes) of immune-related genes upon TA+β infection or whitefly infestation were compared with those in untreated Col-0 plants. (**D**) The expression of genes associated with plant immune responses in TA+β-infected *Arabidopsis* without or with whitefly infestation. TA+β-inoculated plants at 14dpi were infested with whiteflies for 6 h. Relative expression levels (fold changes) of immune-related genes upon whitefly infestation were compared with those in untreated TA+β-inoculated Col-0 plants (Mock). Bars represent means ± SD (n=3) (*, P< 0.05; **, P< 0.01; Student’s *t*-tests).

**Figure 1-figure supplement 2.**
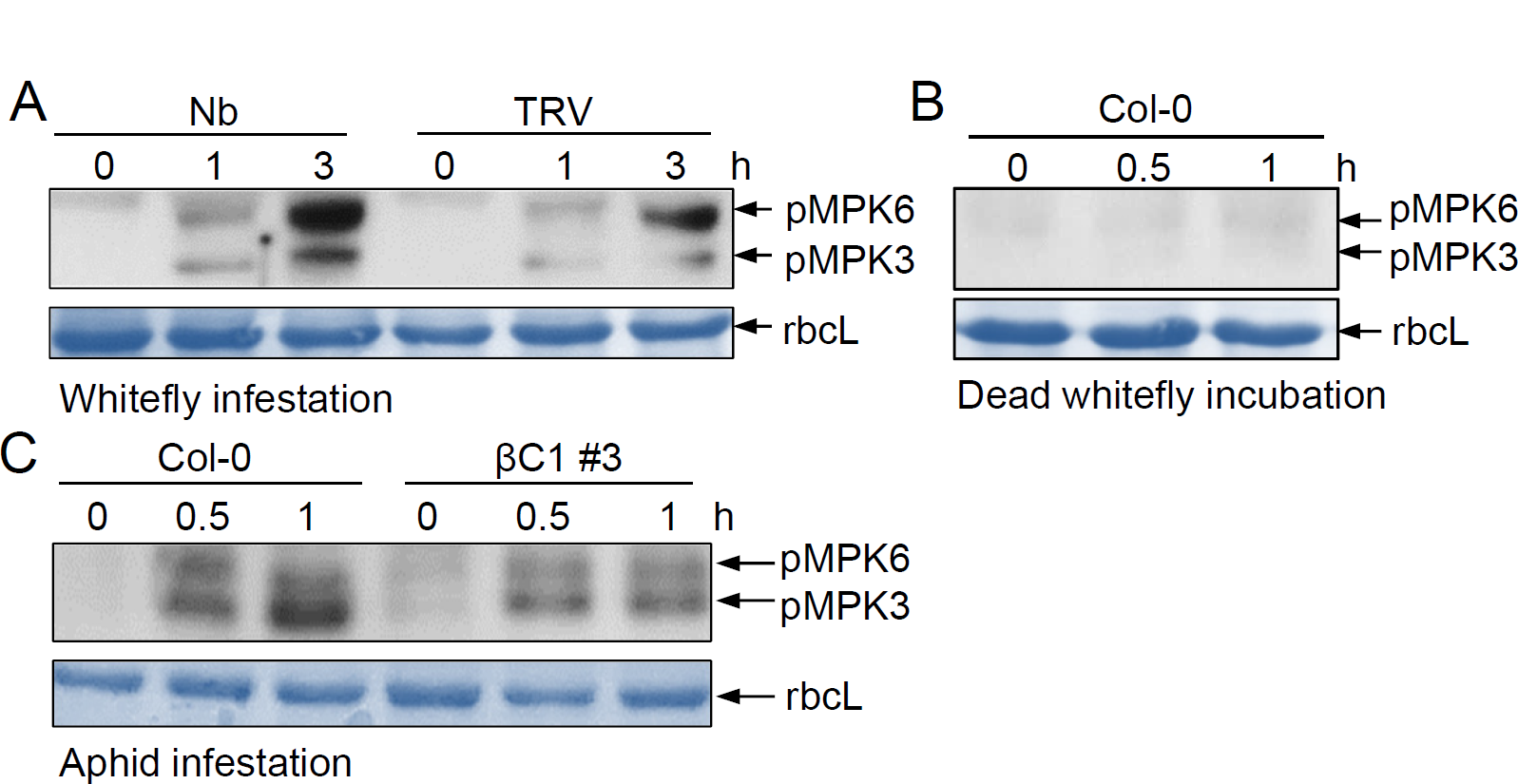
Herbivore-induced defense suppression fails in plants infected by Tobacco rattle virus (TRV) or aphids. (**A**) The effects of MPK3/MPK6 activation by TRV infection upon whitefly infestation in Nb plants. Nb plants were inoculated with TRV. At 14 dpi, plants were infested with whiteflies for the indicated periods. (**B**) The effects of MPK3/MPK6 activation by βC1 protein coupled with aphid infestation in *Arabidopsis*. Plants were infestated with 30 aphids for the indicated periods. (**C**) Dead whiteflies fail to activate plant MAPKs. Cold treated dead whiteflies were applied on Col-0 leaf surface. The phosphorylation levels of plant MPK3 and MPK6 were determined by immunoblot analysis using anti-phospho-p44/42 antibody. Stained membrane bands of the large subunit of Rubisco (rbcL) were used as a loading control.

**Figure 1-figure supplement 3.**
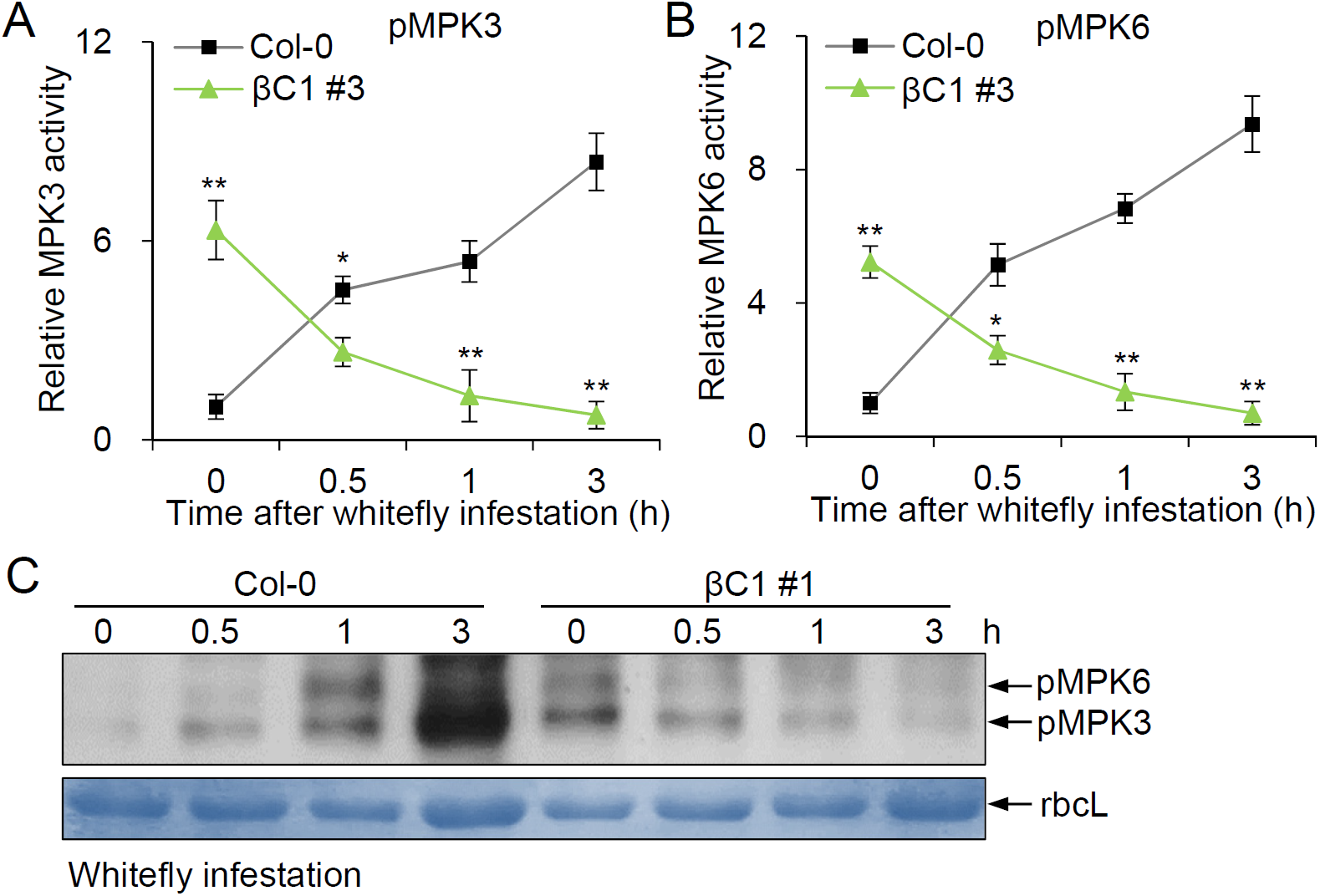
Herbivore-induced defense suppression requires βC1 protein in plants. (**A, B**) Quantitative analysis on activities of MPK3 (**A**) and MPK6 (**B**) in *Arabidopsis* plants upon whitefly infestation. MAPK activity was analysied by ImageJ software to quantify band intensities from the immunoblot analysis in Figure 1C. Bars represent means ± SD (n=3) (*, P< 0.05; **, P< 0.01; Student’s *t*-tests). (**C**) The effects of MPK3/MPK6 activation by βC1 protein upon whitefly infestation in *Arabidopsis*. Plants were infested with 50 whiteflies for the indicated periods. The phosphorylation levels of *Arabidopsis* MPK3 and MPK6 were determined by immunoblot analysis using anti-phospho-p44/42 antibody.

**Figure 2-figure supplement 1.**
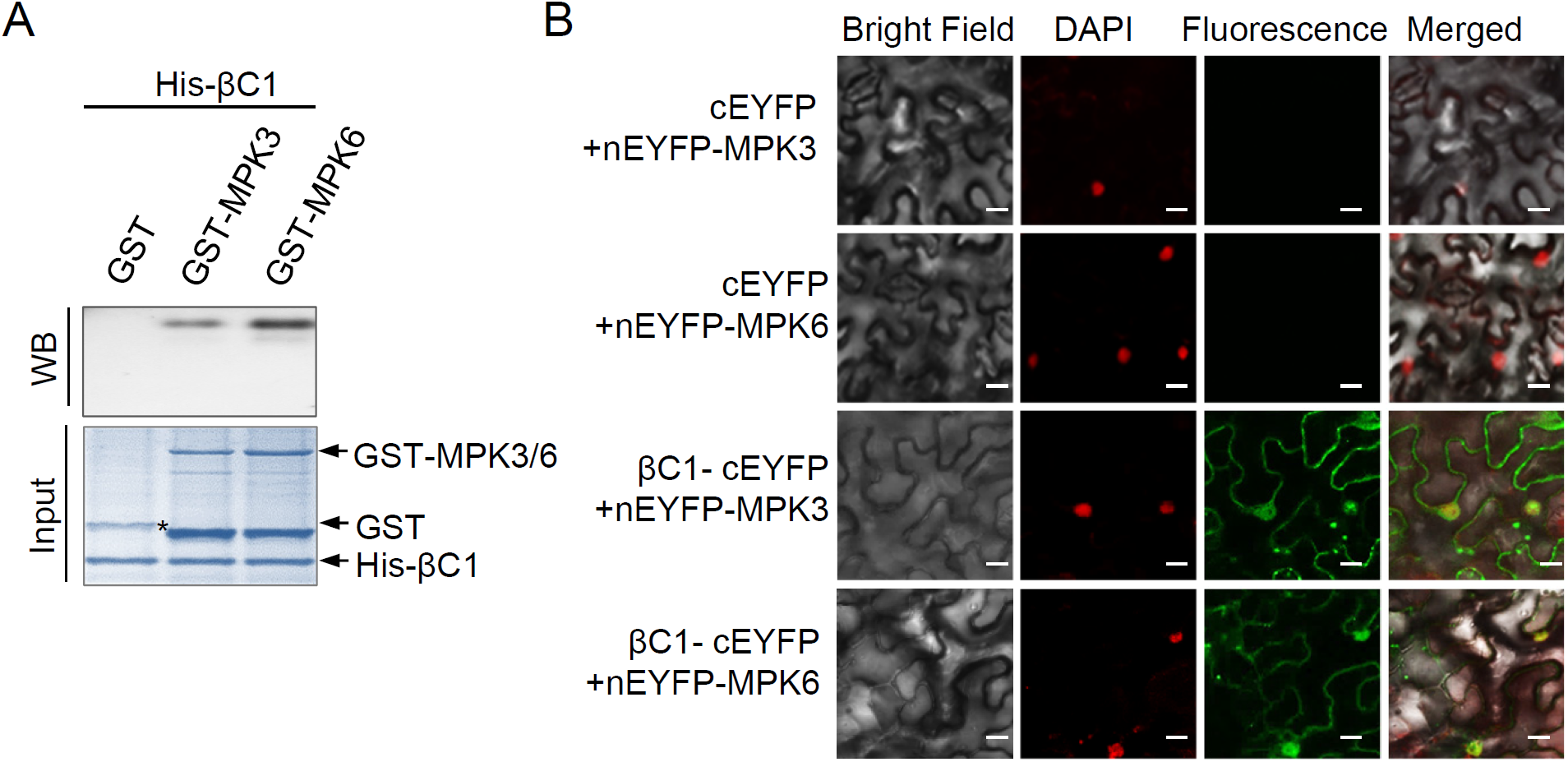
βC1 interacts with *Arabidopsis* MPK3 and MPK6. (**A**) *In vitro* pull-down assay. Two micrograms of His-βC1 fusion protein was used to pull-down 2 μg GST or GST fusion protein. Immunoblot analysis was performed using anti-GST antibody to detect the associated proteins. Membranes were stained with Coomassie brilliant blue to monitor the input protein amounts. (**B**) BiFC analysis shows interaction between βC1 and MPK3 or MPK6. Fluorescence was observed when βC1 fused with the C-terminal part of EYFP complemented MPK3 or MPK6 fused with the N-terminal part of EYFP. Nuclei of tobacco leaf epidermal cells were stained with DAPI. Unfused cEYFP was used as a negative control. Scale bars = 50 μm.

**Figure 2-figure supplement 2.**
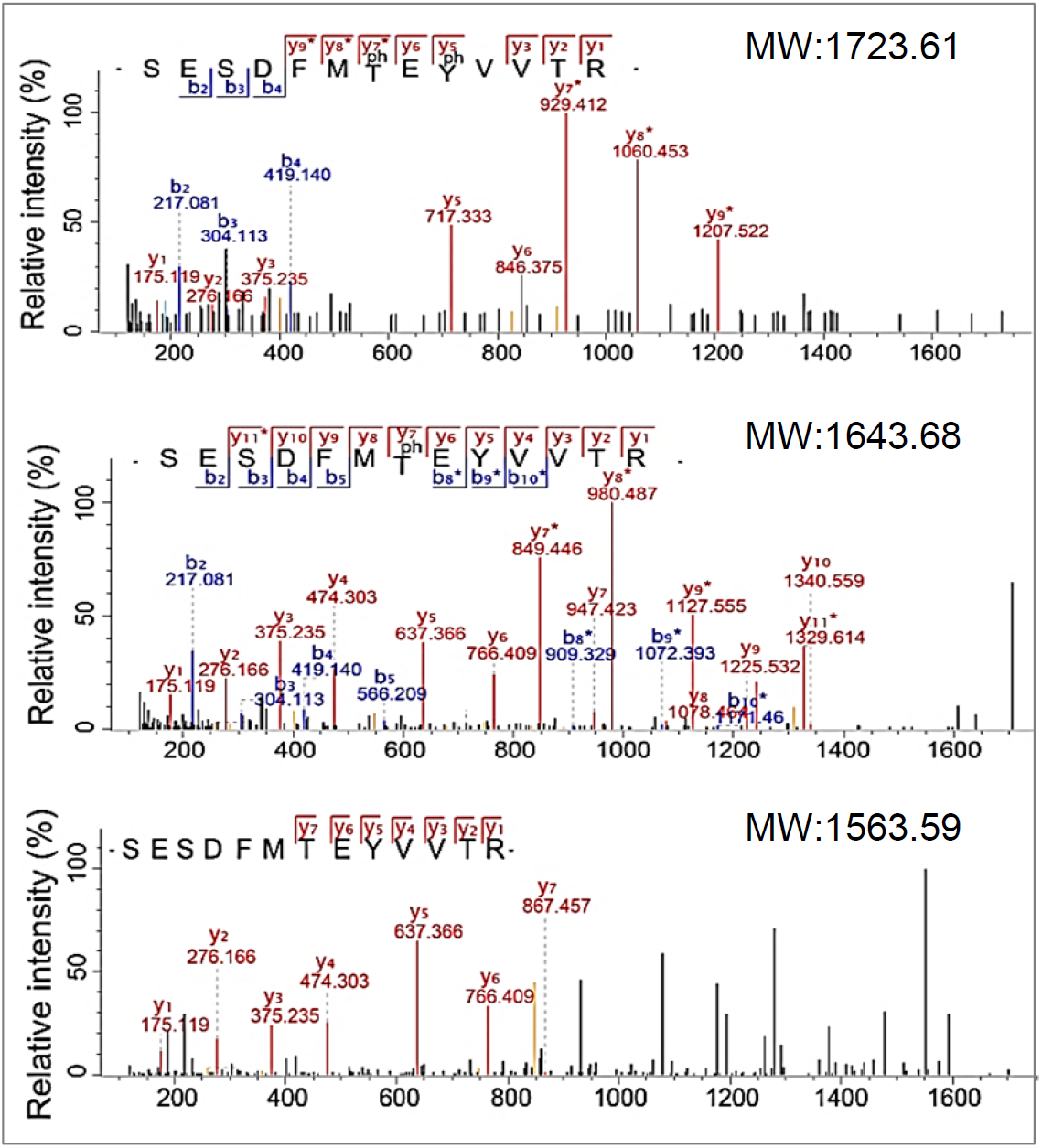
βC1 is a protein phosphatase. The βC1-treated MPK6 phosphopeptide was analyzed by tandem mass spectrometry. The fragmentation profile was indicated by the “b” and “y” ions. The “ph” in the indicated peptide sequence denotes the phosphorylated sites. The “*” indicated the peaks after the neutral loses a phosphate group.

**Figure 2-figure supplement 3.**
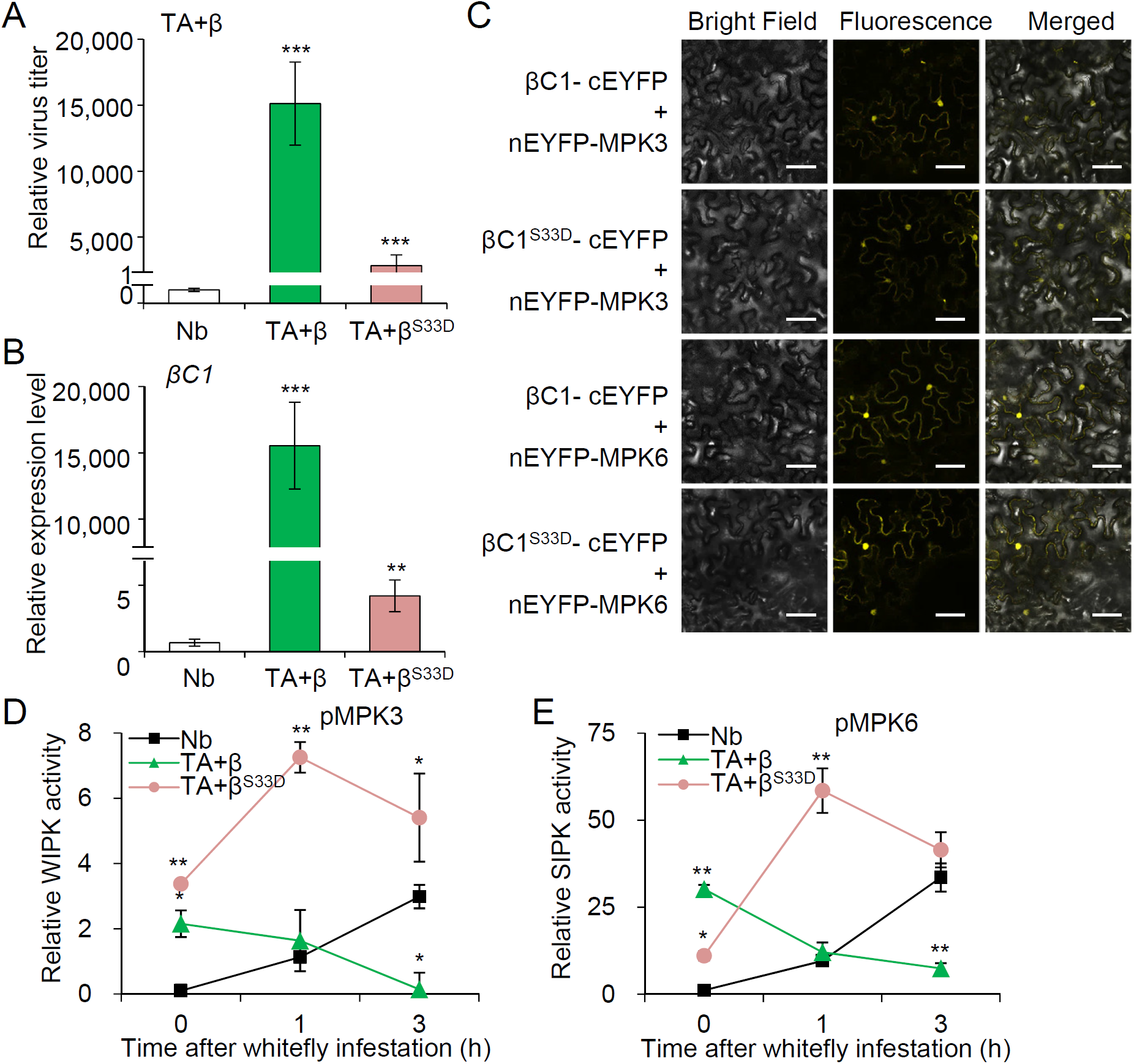
Herbivore-induced defense suppression fails in a phosphorylation mimic of βC1. (**A**) Viral DNA accumulation was quantified by RT-PCR in healthy Nb plants, and Nb plants infected by TA+β or a phosphate βC1 mutant virus (TA+β^S33D^). Bars represent means ± SD (n=4) (***, P< 0.001; Student’s *t*-tests). (**B**) Relative expression level of βC1 in Nb plants, and Nb plants infected by TA+β or TA+β^S33D^. Bars represent means ± SD (n=4) (**, P< 0.01; ***, P< 0.001; Student’s *t*-tests). (**C**) BiFC analysis shows interactions of wild-type βC1 or the phosphorylation mimic mutant βC1^S33D^ with MPK3/6. Fluorescence was observed when βC1 or βC1^S33D^ fused with the C-terminal part of EYFP complemented MPK3/6 fused with the N-terminal part of EYFP. Scale bars = 50 μm. (**D, E**) Quantitative analysis of MPK3 (**D**) and MPK6 (**E**) activity upon whitefly infestation in healthy Nb plants and Nb plants infected by TA+β, or TA+β^S33D^. MAPK activity was analysied by ImageJ software to quantify band intensities from the immunoblots in Figure 2F. Bars represent means ± SD (n=3) (*, P< 0.05; **, P< 0.01; Student’s *t*-tests).

**Figure 3-figure supplement 1.**
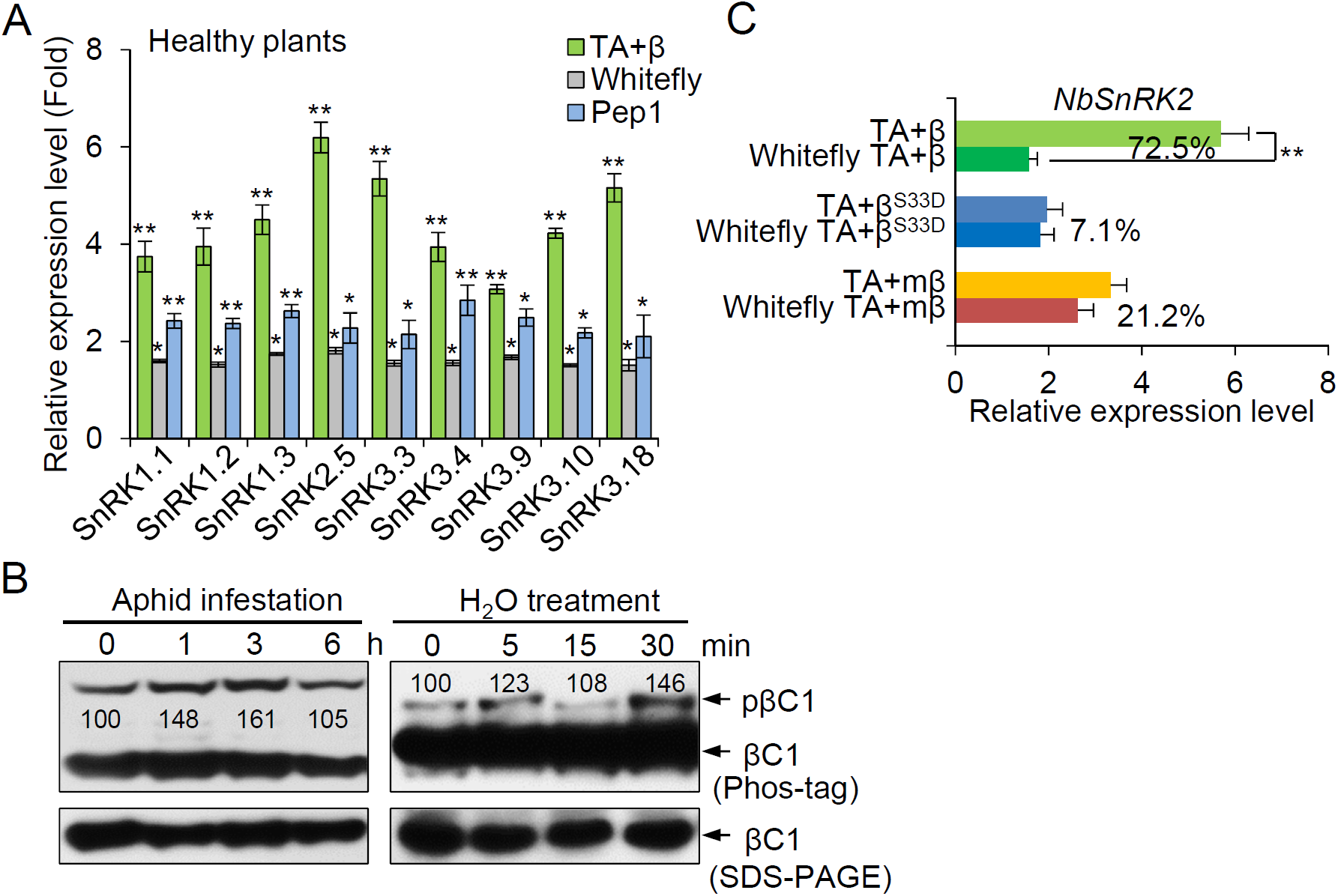
Single stress induces the expression of *SnRKs*. (**A**) Relative expression levels (fold changes) of *Arabidopsis SnRKs* upon each individual treatment (TA+β infection, whitefly infestation, or Pep1 treatment), which were compared with those in healthy Col-0 plants. Bars represent means ± SD (n=3) (*, P< 0.05; **, P< 0.01; Student’s *t*-tests). (**B**) Levels of phosphorylated βC1 protein upon aphid infestation or H_2_O treatment. Phosphorylated pβC1 and unphosphorylated βC1 protein were quantified by band intensities in immunoblots using ImageJ software. The relative level of phosphorylation was normalized against the amount of each treatment at 0 h and shown below each band. (**C**) Relative expression levels of *NbSnRK2* in Nb plants infected by TA+β, TA+β^S33D^, or TA+mβ without or with whitefly infestation for 6 h. Bars represent means ± SD (n=3) (**, P< 0.01; Student’s *t*-tests).

**Figure 5-figure supplement 1.**
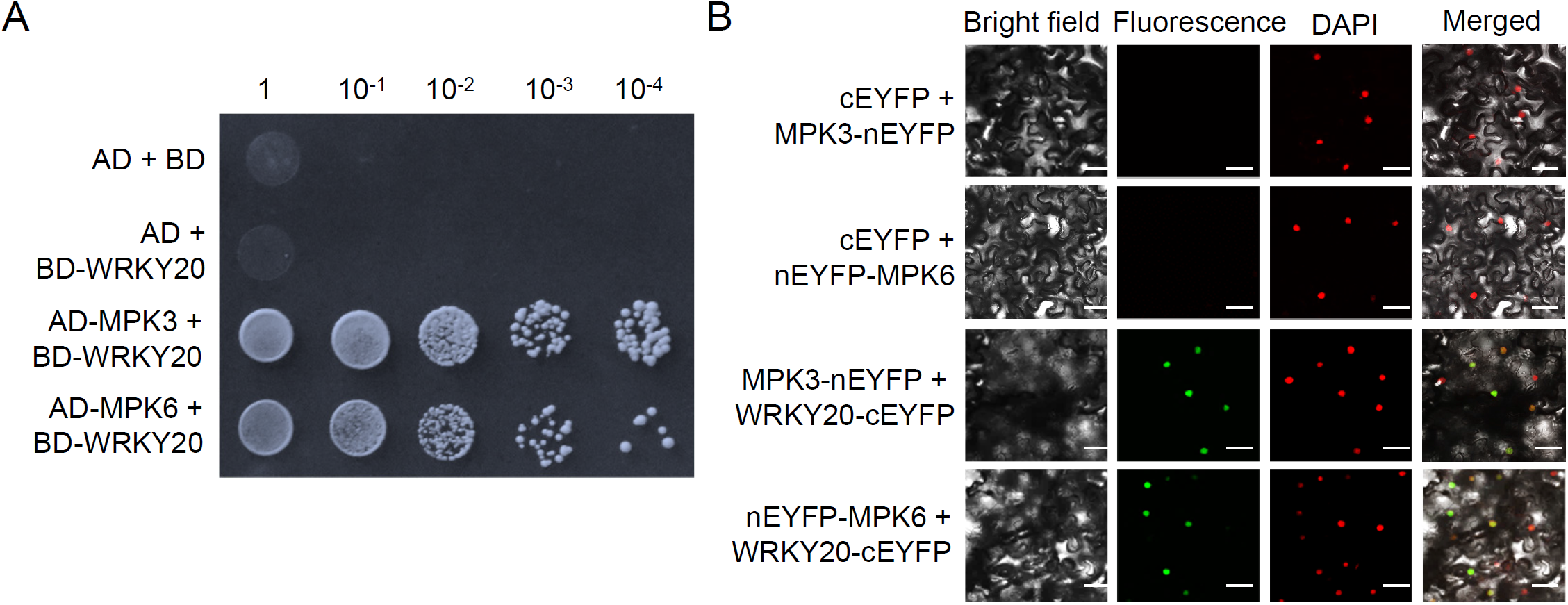
WRKY20 interacts with *Arabidopsis* MPK3/6. (**A**) Interaction of MPK3 or MPK6 with WRKY20 in a yeast two-hybrid system. (**B**) BiFC analysis of MPK3 or MPK6 interaction with WRKY20. Unfused cEYFP was used as a negative control. Scale bars= 50 μm.

**Figure 5-figure supplement 2.**
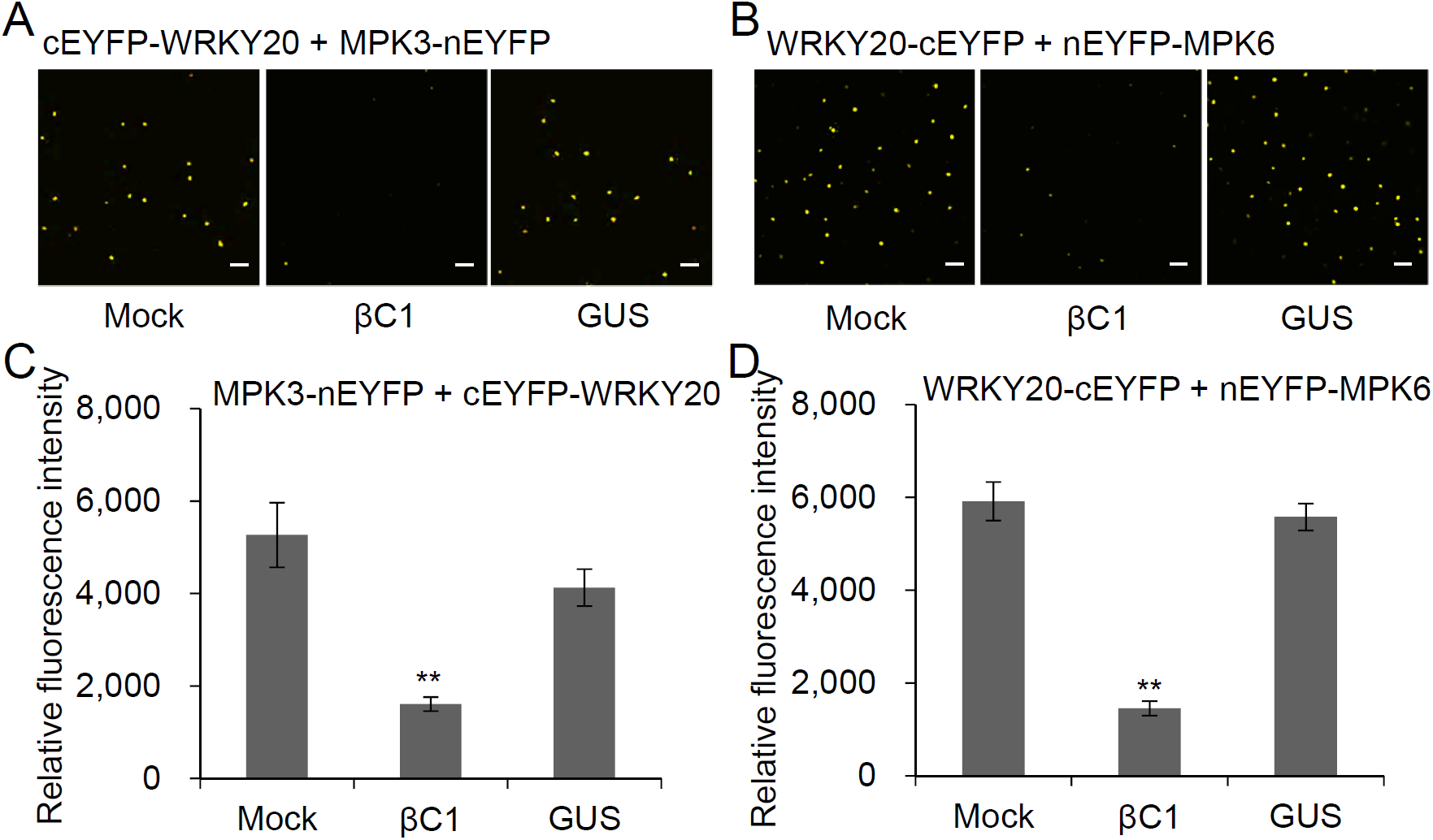
βC1 interferes with the interactions between WRKY20 and MPK3 or MPK6. (**A, B**) Modified BiFC assay was performed to test effects of βC1 on the interaction of WRKY20 with MPK3 (**A**) or MPK6 (**B**). Scale bars= 50 μm. (**C**,**D)** Quantitative data of EYFP fluorescence intensity shows effects of βC1 on the interaction of WRKY20 with MPK3 (**C**) or MPK6 (**D**). Bars represent means ± SD (n=20) (**, P< 0.01; Student’s *t*-tests).

**Figure 5-figure supplement 3.**
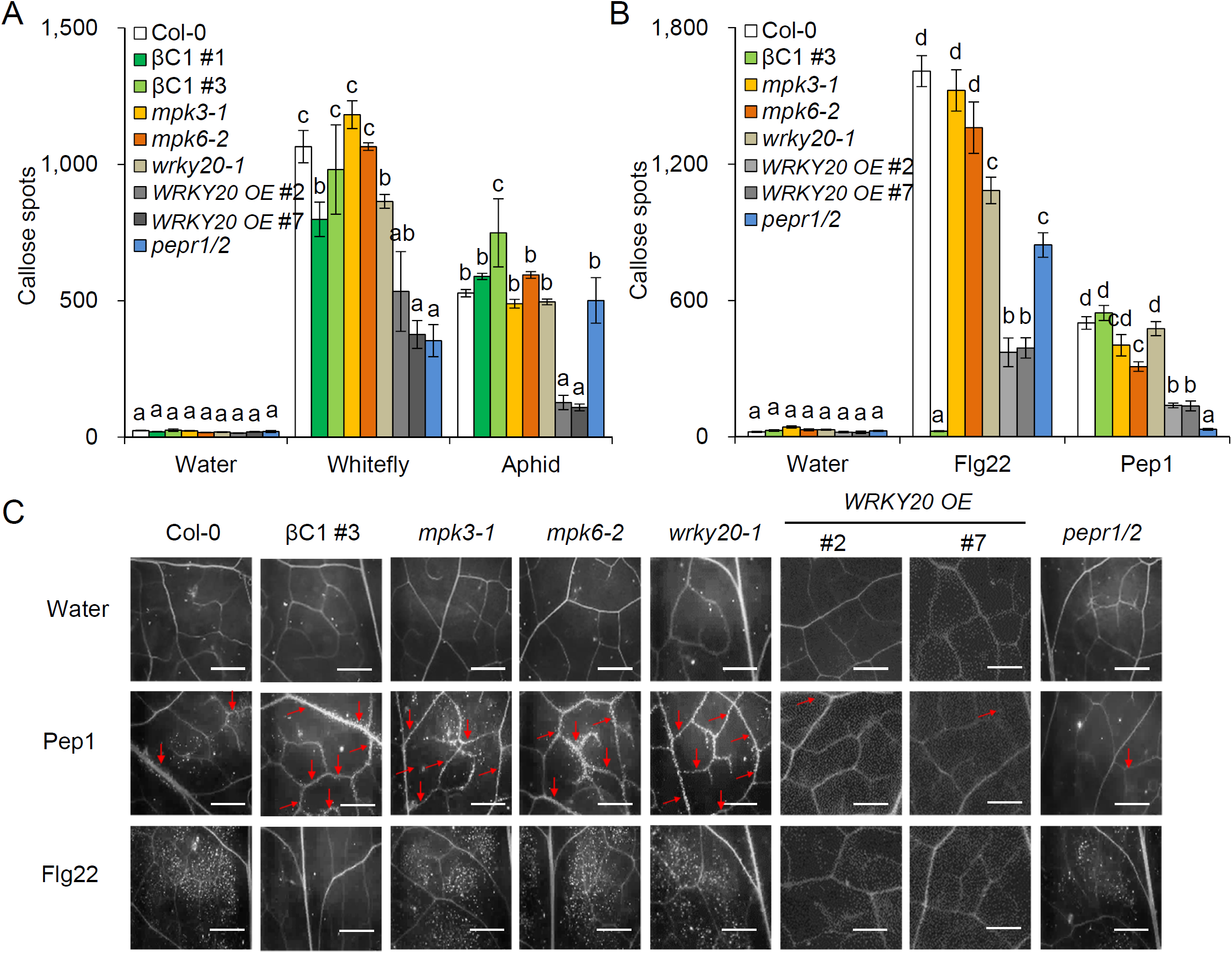
*Arabidopsis* WRKY20 suppresses herbivore-induced callose response. (**A**) The number of callose spots by treated with water, whiteflies or aphids in *Arabidopsis* leaves was counted on each leaf using ImageJ software. Bars represent means ± SD (n=15). (**B**) The number of callose spots by treated with water, 1 μM Pep1 or 1 μM Flg22 for 12 hours in *Arabidopsis* leaves was counted on each leaf using ImageJ software. Bars represent means ± SD (n=15). (**C**) Representative images of callose deposition in *Arabidopsis* leaves upon water, 1 μM Pep1 or 1 μM Flg22 treatment for 12 hours. Scale bars = 300 μm. Lowercase letters indicate significant differences among different lines according to one-way ANOVA followed by Duncan’s multiple range test (P < 0.05).

**Figure 6-figure supplement 1.**
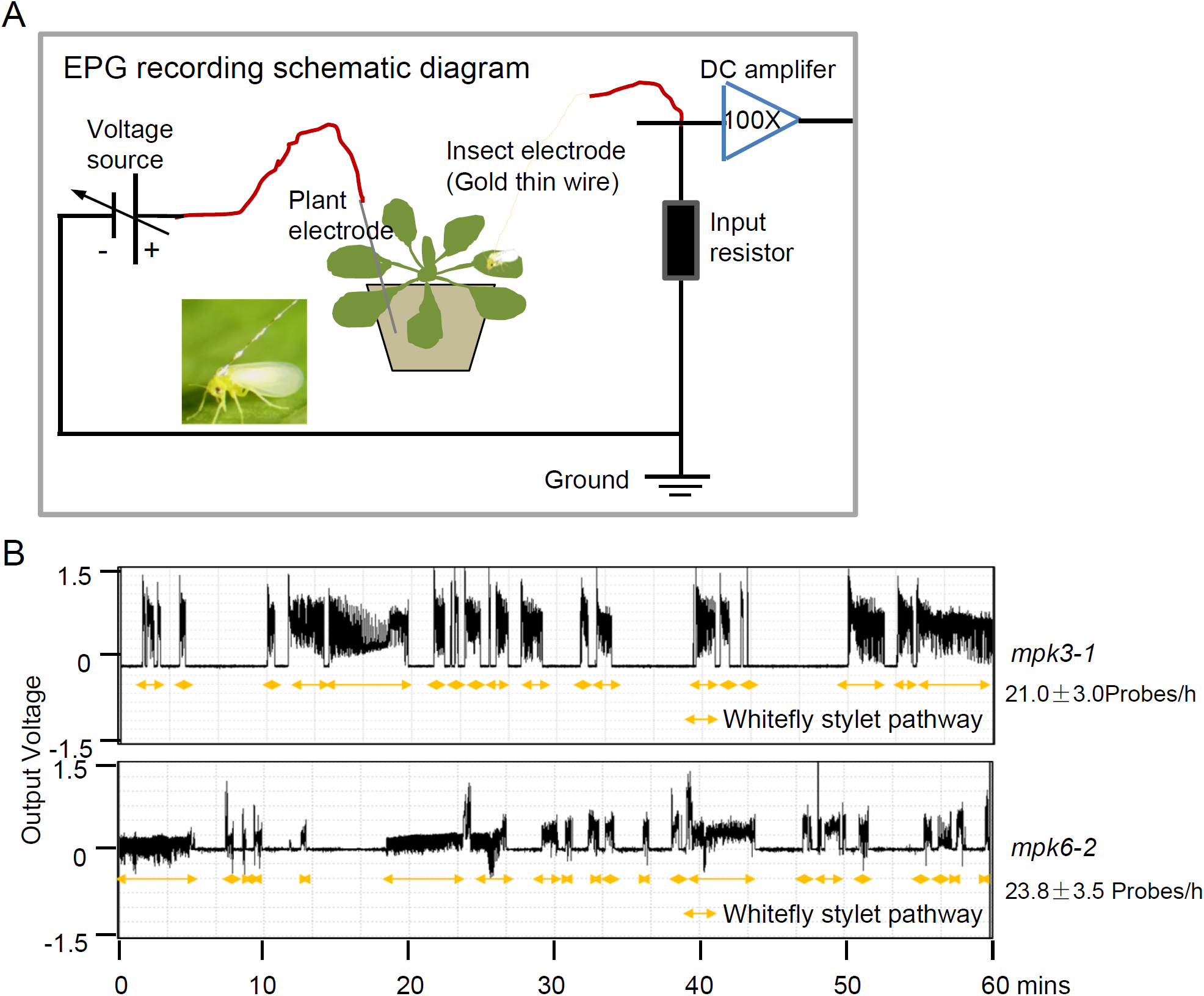
Whitefly probing frequency is improved in *MPK3* or *MPK6* deficiency *Arabidopsis*. (**A**) The electrical penetration graph (EPG) recording schematic diagram. The output probe from the monitor was placed in the soil of the potted plant. The input probe with a thin gold wire (2 cm in length and 12.7 μm in diameter) from the monitor was attached to the pronotum of each whitefly by using a small droplet of silver glue. The EPGs were recorded from insects and plants that were enclosed in a Faraday cage. Output from both monitors was recorded using the Giga-8 EPG system and Stylet+d (EPG Systems) software on a computer. (**B**) Representative EPG waveforms of whitefly stylet pathway phases on two *mpk* mutant *Arabidopsis* plants. Stylet pathways for whitefly represent stylet movement in the intercellular apoplastic space.

## References

Adam L, Somerville SC. 1996. Genetic characterization of five powdery mildew disease resistance loci in *Arabidopsis thaliana*. The Plant Journal 9:341–356. DOI: 10.1046/j.1365-313X.1996.09030341.x, PMID: 8919911

Bak A, Cheung AL, Yang C, Whitham SA, Casteel CL. 2017. A viral protease relocalizes in the presence of the vector to promote vector performance. Nature Communications 8:14493. DOI: 10.1038/ncomms14493, PMID: 28205516

Bartels S, Lori M, Mbengue M, van Verk M, Klauser D, Hander T, Böni R, Robatzek S, Boller T. 2013. The family of Peps and their precursors in Arabidopsis: differential expression and localization but similar induction of pattern-triggered immune responses. Journal of Experimental Botany 64:5309–5321. DOI: 10.1093/jxb/ert330, PMID: 24151300

Chen X, Cao R, Zhong W. 2019. Host calcium channels and pumps in viral infections. Cells 9:94. DOI: 10.3390/cells9010094, PMID: 31905994

Choi HW, Klessig DF. 2016. DAMPs, MAMPs, and NAMPs in plant innate immunity. BMC Plant Biology 16:232. DOI: 10.1186/s12870-016-0921-2, PMID: 27782807

Clay NK, Adio AM, Denoux C, Jander G, Ausubel FM. 2009. Glucosinolate metabolites required for an *Arabidopsis* innate immune response. Science 323:95–101. DOI: 10.1126/science.1164627, PMID: 19095898

Eigenbrode SD, Bosque-Pérez NA, Davis TS. 2018. Insect-borne plant pathogens and their vectors: ecology, evolution, and complex interactions. Annual Review of Entomology 63:169–191. DOI: 10.1146/annurev-ento-020117-043119, PMID: 28968147

Erb M, Reymond P. 2019. Molecular Interactions Between Plants and Insect Herbivores. Annual Review of Plant Biology 70:527–557. DOI: 10.1146/annurev-arplant-050718-095910, PMID: 30786233

Flury P, Klauser D, Schulze B, Boller T, Bartels S. 2013. The anticipation of danger: microbe-associated molecular pattern perception enhances AtPep-triggered oxidative burst. Plant Physiology 161:2023–2035. DOI: 10.1104/pp.113.216077, PMID: 23400703

Fridborg I, Grainger J, Page A, Coleman M, Findlay K, Angell S. 2003. TIP, a novel host factor linking callose degradation with the cell-to-cell movement of Potato virus X. Molecular Plant Microbe Interactions 16:132–140. DOI: 10.1094/MPMI.2003.16.2.132, PMID: 12575747

Fu ZQ, Dong XN. 2013. Systemic acquired resistance: turning local infection into global defense. Annual Review of Plant Biology 64:839–863. DOI: 10.1146/annurev-arplant-042811-105606, PMID: 23373699

García-Sastre A. 2017. Ten strategies of interferon evasion by viruses. Cell Host Microbe 22:176–184. DOI: 10.1016/j.chom.2017.07.012, PMID: 28799903

Gnanasekaran P, KishoreKumar R, Bhattacharyya D, Vinoth Kumar R, Chakraborty S. 2019. Multifaceted role of geminivirus associated betasatellite in pathogenesis. Molecular Plant Pathology 20:1019–1033. DOI: 10.1111/mpp.12800, PMID: 31210029

Gomez-Gomez L, Felix G, Boller T. 1999. A single locus determines sensitivity to bacterial flagellin in *Arabidopsis thaliana*. Plant Journal 18:277–284. DOI: 10.1046/j.1365-313x.1999.00451.x, PMID: 10377993

Guo H, Gu L, Liu F, Chen F, Ge F, Sun Y. 2018. Aphid-borne viral spread is enhanced by virus-induced accumulation of plant reactive oxygen species. Plant Physiology 179:143–155. DOI: 10.1104/pp.18.00437, PMID: 30381318

Haxim Y, Ismayil A, Jia Q, Wang Y, Zheng XY, Chen TY, Qian LC, Liu N, Wang YJ, Han SJ, Cheng JX, Qi YJ, Hong YG, Liu YL. 2017. Autophagy functions as an antiviral mechanism against geminiviruses in plants. Elife 6:e23897. DOI: 10.7554/eLife.23897, PMID: 28244873

Hazzalin CA, Mahadevan LC. 2002. MAPK-regulated transcription: a continuously variable gene switch? Nature Reviews Molecular Cell Biology. 3:30–40. DOI: 10.1038/Nrm715, PMID: 11823796

He WB, Li J, Liu SS. 2015. Differential profiles of direct and indirect modification of vector feeding behaviour by a plant virus. Scientific Reports 5:7682. DOI: 10.1038/Srep07682, PMID: 25567524

Hooks CR, Fereres A. 2006. Protecting crops from non-persistently aphid-transmitted viruses: a review on the use of barrier plants as a management tool. Virus Research 120:1–16. DOI: 10.1016/j.virusres.2006.02.006, PMID: 16780985

Huang D, Sun Y, Ma Z, Ke M, Cui Y, Chen Z, Chen C, Ji C, Tran TM, Yang L, Lam SM, Han Y, Shu G, Friml J, Miao Y, Jiang L, Chen X. 2019. Salicylic acid-mediated plasmodesmal closure via Remorin-dependent lipid organization. Proceedings of the National Academy of Sciences of the United States of America 116:21274–21284. DOI: 10.1073/pnas.1911892116, PMID: 31575745

Huffaker A, Pearce G, Ryan CA. 2006. An endogenous peptide signal in *Arabidopsis* activates components of the innate immune response. Proceedings of the National Academy of Sciences of the United States of America 103:10098–10103. DOI: 10.1073/pnas.0603727103, PMID: 16785434

Jia Q, Liu N, Xie K, Dai Y, Han S, Zhao X, Qian L, Wang Y, Zhao J, Gorovits R, Xie D, Hong Y, Liu Y. 2016. CLCuMuB βC1 subverts ubiquitination by interacting with NbSKP1s to enhance geminivirus infection in *Nicotiana benthamiana*. PLoS Pathogens 12:e1005668. DOI: 10.1371/journal.ppat.1005668, PMID: 27315204

Kinoshita E, Kinoshita-Kikuta E, Takiyama K, Koike T. 2006. Phosphate-binding tag, a new tool to visualize phosphorylated proteins. Molecular & Cellular Proteomics 5:749–757. DOI: 10.1074/mcp.T500024-MCP200, PMID: 16340016

Li P, Shu YN, Fu S, Liu YQ, Zhou XP, Liu SS, Wang XW. 2017. Vector and nonvector insect feeding reduces subsequent plant susceptibility to virus transmission. New Phytologist 215:699–710. DOI: 10.1111/nph.14550, PMID: 28382644

Li R, Pei S, Chen B, Song Y, Zhang T, Yang W, Shaman J. 2020. Substantial undocumented infection facilitates the rapid dissemination of novel coronavirus (SARS-CoV2). Science 368:489–493. DOI: 10.1126/science.abb3221, PMID: 32179701

Li R, Weldegergis BT, Li J, Jung C, Qu J, Sun Y, Qian H, Tee C, van Loon JJ, Dicke M, Chua NH, Liu SS, Ye J. 2014. Virulence factors of geminivirus interact with MYC2 to subvert plant resistance and promote vector performance. The Plant Cell 26:4991–5008. DOI: 10.1105/tpc.114.133181, PMID: 25490915

Liu J, Qian C, Cao X. 2016. Post-translational modification control of innate immunity. Immunity 45:15–30. DOI: 10.1016/j.immuni.2016.06.020, PMID: 27438764

Liu SS, De Barro PJ, Xu J, Luan JB, Zang LS, Ruan YM, Wan FH. 2007. Asymmetric mating interactions drive widespread invasion and displacement in a whitefly. Science 318:1769–1772. DOI: 10.1126/science.1149887, PMID: 17991828

Liu Y, Liu J, Du S, Shan C, Nie K, Zhang R, Li XF, Zhang R, Wang T, Qin CF, Wang P, Shi PY, Cheng G. 2017. Evolutionary enhancement of Zika virus infectivity in *Aedes aegypti* mosquitoes. Nature 545:482–486. DOI: 10.1038/nature22365, PMID: 28514450

Liu Y, Zhang S. 2004. Phosphorylation of 1-aminocyclopropane-1-carboxylic acid synthase by MPK6, a stress-responsive mitogen-activated protein kinase, induces ethylene biosynthesis in Arabidopsis. The Plant Cell 16:3386–3399. DOI: 10.1105/tpc.104.026609, PMID: 15539472

Liu ZX, Wu Y, Yang F, Zhang YY, Chen S, Xie Q, Tian XJ, Zhou JM. 2013. BIK1 interacts with PEPRs to mediate ethylene-induced immunity. Proceedings of the National Academy of Sciences of the United States of America 110:6205–6210. DOI: 10.1073/pnas.1215543110, PMID: 23431184

Luan JB, Yao DM, Zhang T, Walling LL, Yang M, Wang YJ, Liu SS. 2013. Suppression of terpenoid synthesis in plants by a virus promotes its mutualism with vectors. Ecology Letters 16:390–398. DOI: 10.1111/ele.12055, PMID: 23279824

Luna E, Pastor V, Robert J, Flors V, Mauch-Mani B, Ton J. 2011. Callose deposition: a multifaceted plant defense response. Molecular Plant Microbe Interactions 24:183–193. DOI: 10.1094/MPMI-07-10-0149, PMID: 20955078

Martiniere A, Bak A, Macia JL, Lautredou N, Gargani D, Doumayrou J, Garzo E, Moreno A, Fereres A, Blanc S, Drucker M. 2013. A virus responds instantly to the presence of the vector on the host and forms transmission morphs. Elife 2:e00183. DOI: 10.7554/eLife.00183, PMID: 23358702

Mauck KE, Kenney J, Chesnais Q. 2019. Progress and challenges in identifying molecular mechanisms underlying host and vector manipulation by plant viruses. Current Opinion in Insect Science 33:30082–30088. DOI: 10.7554/eLife.00183, PMID: 31358199

Prado Maluta NK, Garzo E, Moreno A, Navas-Castillo J, Fiallo-Olive E, Spotti Lopes JR, Fereres A. 2017. Stylet penetration activities of the whitefly *Bemisia tabaci* associated with inoculation of the crinivirus *Tomato chlorosis virus*. Journal of General Virology 98:1515–1520. DOI: 10.1099/jgv.0.000783, PMID: 28613151

Qu J, Ye J, Geng YF, Sun YW, Gao SQ, Zhang BP, Chen W, Chua NH. 2012. Dissecting functions of *KATANIN* and *WRINKLED1* in cotton fiber development by virus-induced gene silencing. Plant Physiology 160:738–748. DOI: 10.1104/pp.112.198564, PMID: 22837356

Saijo Y, Loo EP, Yasuda S. 2018. Pattern recognition receptors and signaling in plant-microbe interactions. Plant Journal 93:592–613. DOI: 10.1111/tpj.13808, PMID: 29266555

Schuman MC, Baldwin IT. 2016. The layers of plant responses to insect herbivores. Annual Review of Entomology 61:373–394. DOI: 10.1146/annurev-ento-010715-023851, PMID: 26651543

Sethi V, Raghuram B, Sinha AK, Chattopadhyay S. 2014. A mitogen-activated protein kinase cascade module, MKK3-MPK6 and MYC2, is involved in blue light-mediated seedling development in Arabidopsis. The Plant Cell 26:3343–3357. DOI: 10.1105/tpc.114.128702, PMID: 25139007

Sharrocks AD, Yang SH, Galanis A. 2000. Docking domains and substrate-specificity determination for MAP kinases. Trends in Biochemical Sciences 25:448–453. DOI: 10.1016/s0968-0004(00)01627-3, PMID: 10973059

Shen Q, Liu Z, Song F, Xie Q, Hanley-Bowdoin L, Zhou X. 2011. Tomato SlSnRK1 protein interacts with and phosphorylates βC1, a pathogenesis protein encoded by a geminivirus β-satellite. Plant Physiology 157:1394–1406. DOI: 10.1104/pp.111.184648, PMID: 21885668

Stafford CA, Walker GP, Ullman DE. 2011. Infection with a plant virus modifies vector feeding behavior. Proceedings of the National Academy of Sciences of the United States of America 108:9350–9355. DOI: 10.1073/pnas.1100773108, PMID: 21606372

Tsuda K, Somssich IE. 2015. Transcriptional networks in plant immunity. New Phytologist 206:932–947. DOI: 10.1111/nph.13286, PMID: 25623163

Vasilakis N, Tesh RB. 2015. Insect-specific viruses and their potential impact on arbovirus transmission. Current Opinion in Virology 15:69–74. DOI: 10.1016/j.coviro.2015.08.007, PMID: 26322695

Vincent TR, Avramova M, Canham J, Higgins P, Bilkey N, Mugford ST, Pitino M, Toyota M, Gilroy S, Miller AJ, Hogenhout SA, Sanders D. 2017. Interplay of plasma membrane and vacuolar ion channels, together with BAK1, elicits rapid cytosolic calcium elevations in Arabidopsis during aphid feeding. The Plant Cell 29:1460–1479. DOI: 10.1105/tpc.17.00136, PMID: 28559475

Wang N, Zhao P, Ma Y, Yao X, Sun Y, Huang X, Jin J, Zhang Y, Zhu C, Fang R, Ye J. 2019. A whitefly effector Bsp9 targets host immunity regulator WRKY33 to promote performance. Philosophical Transactions of The Royal Society B-Biological Sciences 374:20180313. DOI: 10.1098/rstb.2018.0313, PMID: 30967015

Weaver SC, Charlier C, Vasilakis N, Lecuit M. 2018. Zika, chikungunya, and other emerging vector-borne viral diseases. Annual Review of Medicine 69:395–408. DOI: 10.1146/annurev-med-050715-105122, PMID: 28846489

Wu X, Xu S, Zhao P, Zhang X, Yao X, Sun Y, Fang R, Ye J. 2019. The Orthotospovirus nonstructural protein NSs suppresses plant MYC-regulated jasmonate signaling leading to enhanced vector attraction and performance. PLoS Pathogens 15:e1007897. DOI: 10.1371/journal.ppat.1007897, PMID: 31206553

Wu X, Ye J. 2020. Manipulation of Jasmonate Signaling by Plant Viruses and Their Insect Vectors. Viruses 12. DOI: 10.3390/v12020148, PMID: 32012772

Wurzinger B, Nukarinen E, Nagele T, Weckwerth W, Teige M. 2018. The SnRK1 kinase as central mediator of energy signaling between different organelles. Plant Physiology 176:1085–1094. DOI: 10.1104/pp.17.01404, PMID: 29311271

Xu J, Meng J, Meng X, Zhao Y, Liu J, Sun T, Liu Y, Wang Q, Zhang S. 2016. Pathogen-responsive MPK3 and MPK6 reprogram the biosynthesis of indole glucosinolates and their derivatives in *Arabidopsis* immunity. The Plant Cell 28:1144–1162. DOI: 10.1105/tpc.15.00871, PMID: 27081184

Yang JY, Iwasaki M, Machida C, Machida Y, Zhou X, Chua NH. 2008. βC1, the pathogenicity factor of TYLCCNV, interacts with AS1 to alter leaf development and suppress selective jasmonic acid responses. Genes & Development 22:2564–2577. DOI: 10.1101/gad.1682208, PMID: 18794352

Yang X, Guo W, Li F, Sunter G, Zhou X. 2019. Geminivirus-associated betasatellites: Exploiting chinks in the antiviral arsenal of plants. Trends in Plant Science 24:519–529. DOI: 10.1016/j.tplants.2019.03.010, PMID: 31003895

Ye J, Yang JY, Sun YW, Zhao PZ, Gao SQ, Jung C, Qu J, Fang RX, Chua NH. 2015. Geminivirus activates ASYMMETRIC LEAVES 2 to accelerate cytoplasmic DCP2-mediated mRNA turnover and weakens RNA silencing in *Arabidopsis*. PLoS Pathogens 11:e1005196. DOI: 10.1371/journal.ppat.1005196, PMID: 26431425

Zavaliev R, Ueki S, Epel BL, Citovsky V. 2011. Biology of callose (β-1,3-glucan) turnover at plasmodesmata. Protoplasma 248:117–130. DOI: 10.1007/s00709-010-0247-0, PMID: 21116665

Zhang J, Shao F, Li Y, Cui H, Chen L, Li H, Zou Y, Long C, Lan L, Chai J, Chen S, Tang X, Zhou JM. 2007. A *Pseudomonas syringae* effector inactivates MAPKs to suppress PAMP-induced immunity in plants. Cell Host Microbe 1:175–185. DOI: 10.1016/j.chom.2007.03.006, PMID: 18005697

Zhang T, Luan JB, Qi JF, Huang CJ, Li M, Zhou XP, Liu SS. 2012. Begomovirus-whitefly mutualism is achieved through repression of plant defences by a virus pathogenicity factor. Molecular Ecology 21:1294–1304. DOI: 10.1111/j.1365-294X.2012.05457.x, PMID: 22269032

Zhao P, Yao X, Cai C, Li R, Du J, Sun Y, Wang M, Zou Z, Wang Q, Kliebenstein DJ, Liu SS, Fang RX, Ye J. 2019. Viruses mobilize plant immunity to deter nonvector insect herbivores. Science Advances 5:eaav9801. DOI: 10.1126/sciadv.aav9801, PMID: 31457079

Zhong X, Wang ZQ, Xiao R, Cao L, Wang Y, Xie Y, Zhou X. 2017. Mimic phosphorylation of a βC1 protein encoded by TYLCCNB impairs its functions as a viral suppressor of RNA silencing and a symptom determinant. Journal of Virology 91:e00300–00317. DOI: 10.1128/JVI.00300-17, PMID: 28539450

Zhou XP. 2013. Advances in understanding begomovirus satellites. Annual Review of Phytopathology 51:357–381. DOI: 10.1146/annurev-phyto-082712-102234, PMID: 23915133

Zhou Z, Zhao Y, Bi G, Liang X, Zhou JM. 2019. Early signalling mechanisms underlying receptor kinase-mediated immunity in plants. Philosophical Transactions of The Royal Society B-Biological Sciences 374:20180310. DOI: 10.1098/rstb.2018.0310, PMID: 30967025

